# scRM: A single-cell foundation model trained on scRNA-seq reads

**DOI:** 10.64898/2026.10.24.761590

**Authors:** Dongze He, Rob Patro, Marcel Nassar, Haiyu Wu, Qi Yan, Glenn Xu, Kristof Torkenczy, Jamin Liu, Stefka Tyanova

## Abstract

Single-cell foundation models currently learn from gene count matrices, which are discretized, species-specific summaries that discard the RNA sequence, splice-site usage, 3^*′*^ UTR isoform identity and the mutational information present in the underlying sequencing reads. Here we introduce single-cell raw data pretraining, a framework that trains foundation models directly on sequencing reads, requiring no read alignment or count matrix, allowing any trained model to be applicable to arbitrary species.

Our first model in this framework, the single-cell read model scRM, pairs two transformer encoders with a hierarchy of attention pools and contains 1.2 million parameters. We trained it as a JEPA on 3,000 mouse neuron cells on a single H100 in one hour, and benchmarked it zero-shot against leading single-cell foundation models with orders of magnitude more parameters and training cells. Our results show that our lightweight model reaches the performance of existing models on all tasks, and exceeds every one of them on perturbation detection tasks whose detection requires sequence-level detail.

To our knowledge, scRM is the first single-cell foundation model trained on sequencing reads, demonstrating that raw data pretraining is not only feasible but competitive at a fraction of the computational cost. We anticipate that scaling this framework and exploring alternative architectures will unlock biological information inaccessible to count models and extend the dataset-specific pretraining it already makes possible. The source code is available at https://github.com/altoslabs/scRM.

## 1 Introduction

Single-cell deep learning models have gone through tremendous developments in recent years, and to date most of them take count matrices as the input. scVI (Lopez et al., 2018) and DCA (Eraslan et al., 2019) fit autoencoders to the count matrix, to denoise it and to represent each cell as a low-dimensional vector that later analysis can use. Other deep models, such as convolutional and graph neural networks and generative adversarial networks, are also trained on the same input and have shown extraordinary successes (Yuan and Bar-Joseph, 2019; Wang et al., 2021a; Marouf et al., 2020). With the successes of transformers and large language models, researchers have also applied them to single-cell data. The first transformer over gene tokens, iSEEEK (Shen et al., 2022), ranked each cell’s genes by expression and fed the ranked gene symbols as a sequence. To use the counts themselves in transformer training, successors turn the count matrix into a corpus a transformer can be trained on, using techniques such as binning, ranking or projecting, because transformers are not designed for count matrices (Cui et al., 2024; Hao et al., 2024; Theodoris et al., 2023).

Count matrices have a rich history in RNA-sequencing data analysis, and they exist because the raw data, the amplified and sequenced cDNA fragments reverse transcribed from RNA, are too big and carry too low an information density for a human analyst to read. The pipeline for generating a count matrix from the raw data mainly consists of two steps, where read alignment aligns observed sequencing reads against the reference genome or transcriptome, and quantification quantifies aligned reads to their gene of origin (Patro et al., 2017; He et al., 2022a). Such count matrices work well for low-resolution protocols like the microarray (Schena et al., 1995) and bulk RNA-seq (Mortazavi et al., 2008), because each sample sums the signal of many cells at once, which gives a dense and information-rich count matrix with the entire population as one sample. For scRNA-seq, however, the resulting count matrices are usually extremely sparse, with up to 90% zeros, which causes difficulties in many downstream analyses (Jiang et al., 2022). This is because the cDNA library must be divided back into thousands or tens of thousands of individual cells, so each sample now sums over one cell rather than over a population. Another defect of relying on count matrices is that, by summing up the sequencing reads to gene counts, all fine-grained information, like the usage of polyadenylation sites, UTR sites, and splicing sites, is discarded along the process, blocking many downstream analysis types (Patrick et al., 2020).

Over time, researchers realized that count matrices alone might not be the optimal input for single-cell foundation model training. On one hand, focusing on gene identity ignores the rich, sequence-level information carried within sequencing reads, such as alternative polyadenylation, 3^*′*^ UTR usage, alternative splicing, and the allele a molecule came from (Patrick et al., 2020; Fansler et al., 2024; Huang and Sanguinetti, 2017; Deng et al., 2014). On the other hand, relying on known gene annotation blocks the trained model from being applied across species, because their gene identities might differ. For example, only 20% of the human genes with a fly ortholog have a strict one-to-one counterpart (Hu et al., 2011). The limitation is hardest on organisms with no assembled genome at all, which are often disease-relevant as viral or bacterial reservoirs and are understudied because the standard tools do not exist for them. Furthermore, not all sequencing reads can be mapped to gene-annotated regions. Wang et al. (2021b) reported that reads from transcriptionally active regions outside annotated genes recover biology that annotation-bound analysis misses, across six species whose annotations are incomplete. Similarly, He and Patro (2024) showed that off-target reads from 10x Chromium 3^*′*^ datasets carry chromatin structure and can improve gene detection.

Because of these drawbacks, researchers explored ways to augment count matrices for single-cell foundation model training. scBERT takes gene identity vectors from a learned embedding, to pass the gene-gene relationship to the model (Yang et al., 2022). UCE represents each gene token with an embedding of that gene’s protein sequence, so training is aware of the sequence behind it (Rosen et al., 2026). scooby adapts Borzoi, which predicts measured coverage from DNA sequence, by freezing its weights and fine-tuning its embeddings (Linder et al., 2025; Hingerl et al., 2025). All these attempts resulted in extraordinary performance, and each adds information the count matrix does not carry, which suggests it alone is too sparse for single-cell foundation model training.

In this project, we propose a novel way to train single-cell foundation models. We bring attention to the very raw scRNA-seq data, the sequencing reads, and use that to build the training corpus. The advantage is that nucleic acid sequences of A, C, G and T are the natural input for language models like transformers, and that they carry the transcribed, active genetic information within the cell, which reflects the actual cellular activity more precisely than passing prebuilt embeddings of genes or proteins. Moreover, once we apply conventional data processing steps like cell barcode correction and UMI deduplication, the deduplicated reads themselves represent the gene activity information.

This type of information is much richer than a count matrix as it carries transcript-level signals, including alternative splicing, 3^*′*^ UTR usage, and every signal listed above. We call this framework **single-cell raw data pretraining**, and the single-cell read model **scRM**, a hierarchical transformer inspired by the byte latent transformer (Pagnoni et al., 2025) and adopted under this setting, is the first model we trained in it (Figure 1, §2).

**Figure 1.**
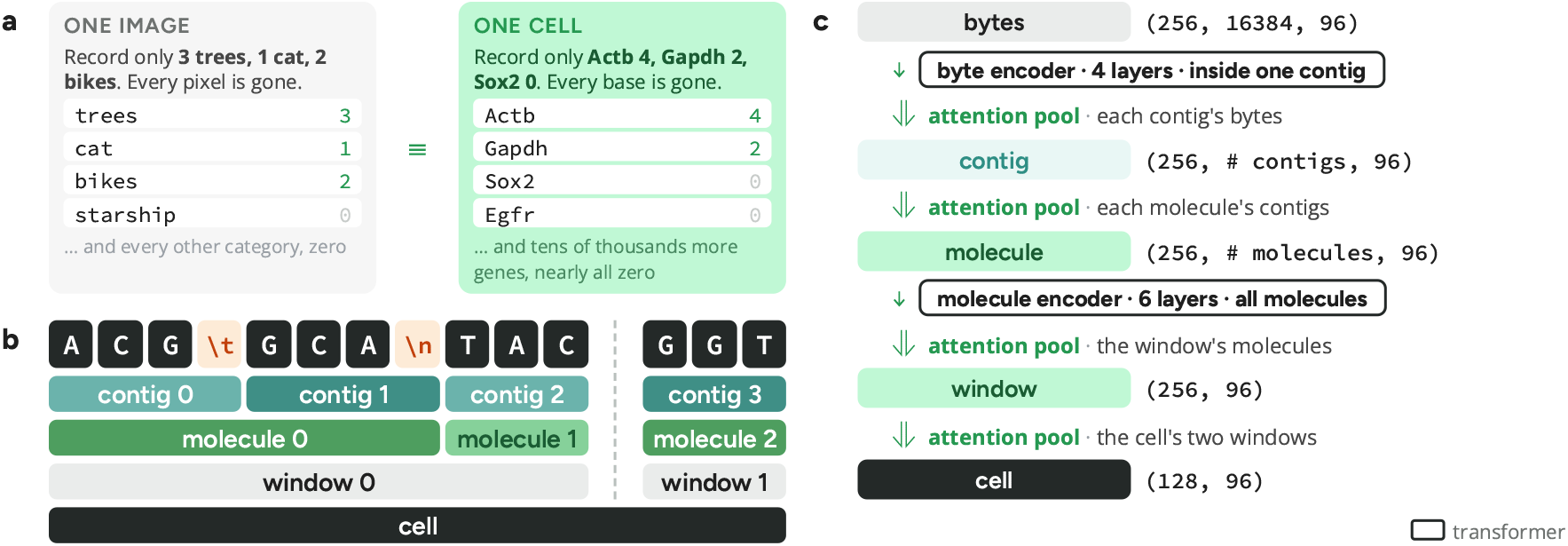
**(a) A count matrix is a feature count table.** Counting the genes in a cell is analogous to counting the objects in an image and discarding every pixel. All existing single-cell foundation models take count matrices as the input. **(b) Information propagation in single-cell raw data pretraining**. Raw sequencing reads are grouped and deduplicated by their cell barcode and UMI. The assembled reads in each UMI are represented by contigs. Disjoint contigs of a UMI are concatenated by a tab (`\t`). UMIs are concatenated by a newline (`\n`). UMIs from the same cell are filled into each input window in random order until the next one would not fit, so the windows partition the cell. Each cell has two windows in a training batch, required by the regularizer loss term. **(c) The scRM model**. scRM takes the constructed input windows directly. It has two transformers, one at the byte level and another at the molecule level. Information is propagated up the hierarchy using attention pools at each of the four boundaries. The shapes noted in each module are one training step (§2.3).

Our contributions come from two folds: Firstly, single-cell raw data pretraining is a highly efficient framework implemented in Rust that trains single-cell foundation models on two things at once, the genetic information a cell’s sequencing reads carry, and the gene expression profile those reads count toward. It uses the rich sequenced reads, rather than read alignment or a count matrix, so models can be trained on one scRNA-seq experiment, and any trained model can be applied to arbitrary species (§2.1). Moreover, scRM shows how little data the framework needs. A 1.2M-parameter model trained on 3,000 mouse neuron cells (i) reaches the leading count foundation models on all five evaluation tasks and predicts cell type in five species from mouse-only training, (ii) exceeds every one of them where predictions require sequence-level detail, and (iii) makes dataset-specific pretraining feasible for the first time, best for species and tissues with little data (§3). Supplementary Material Supp. A gives the related work.

## 2 Method

### 2.1 The single-cell raw data pretraining framework

To build the training corpus from sequencing reads, we provide a highly efficient command-line tool written in Rust, with many modules adopted from alevin-fry (He et al., 2022a). The pipeline takes the sequencing reads and a list of high-confidence cell barcodes from a cell calling tool like QCatch (Gao et al., 2026). As in the conventional pipeline, Read 2, the captured cDNA fragment, builds the training corpus, and Read 1 is parsed for the cell barcode and UMI, identifying the reads from each pre-amplified RNA in each cell. That list is the permit list for cell barcode correction. Only reads labeled by a permitted barcode that occurs in the list are kept. Any other barcode is corrected to a permitted barcode when it is one Hamming distance away from exactly one of them. Otherwise it is discarded. Cell calling is therefore the only place a reference-based tool reaches the corpus build.

The UMIs within a cell have to be deduplicated before their reads are processed, because sequencing errors might write the UMI of one molecule as several similar sequences. We followed alevin-fry to build one directed UMI graph per cell to deduplicate UMIs. Its vertices are the distinct UMI sequences observed in that cell, weighted by its read count. Every vertex is given at most one out-edge, pointing at the neighbor of greatest weight at Hamming distance one, and only when that neighbor is strictly heavier. The rarer sequence is therefore a copy of the more abundant one. No edge is drawn between two neighbors of equal weight. A vertex with no heavier neighbor is a sink. As every edge increases weight, the graph is acyclic with out-degree at most one. Each connected component is therefore a tree whose edges all lead to one sink. We regard each component as a deduplicated UMI, and replace its UMIs with the sink’s. A cell’s molecules carry its gene activity, so a wrong merge takes away a molecule the cell produced and the model reads less activity than it had.

The reads of each deduplicated UMI in each cell are then used to assemble the original RNA sequence associated with the UMI. As the name suggests, one UMI represents one captured mRNA, and its reads are error-prone fragments taken from its poly(A) tail or from an internal adenine repeat, overlapping or disjoint. We reconstruct the captured mRNA sequence by building a de Bruijn graph over the UMI’s reads at *k* = 19. Its vertices are the distinct 19-mers of those reads, each weighted by the number of reads containing it. A directed edge connects two vertices whose 18-mer overlap is observed in a read. The graph is then path-compressed (Marks, 2024), so each vertex spells a maximal non-branching path, i.e., a unitig, represented by the merged 19-mers, weighted by the lowest weight of its 19-mers. We then start a walk at the heaviest vertex not yet used that is not itself one branch of a disagreement, and extend it in both directions, so the best-supported sequence is read off first. The sequence a walk spells is a contig.

Along the walk we solve the branches, the points where two paths disagree, using the Phred score from FASTQ and DADA2’s abundance test (Callahan et al., 2016). Briefly, we ask whether the differences between the branches are errors. A high Phred score on every base of every branch means none of those bases should be wrong, so the branches are just indicating loci sharing short sequences, and each branch that rejoins the other deserves its own contig. A branch that does *not* rejoin is instead consumed into the walk, because its reads have already voted on every column they cover, so emitting it separately would duplicate the contig it diverges from rather than add evidence. Low Phred scores on the mismatched bases suggest the opposite, that the branches are erroneous copies of one molecule, so we merge them and call each mismatched base from the reads covering it using a per-base Phred likelihood model (Fulcrum Genomics, 2026), writing the ambiguous base `N` where that call cannot be made. Two `N`s in a row say the differences are not errors after all, so the contig splits there rather than carrying a run, and `N` always marks a single ambiguous base. No read is discarded for disagreeing with another. One structure is dropped rather than resolved: a homopolymer run of at least *k* bases collapses to a single self-looping vertex, which destroys the run’s length, so that vertex is discarded instead of being spelled at a length the graph cannot know. Contigs on either side of it therefore end *k* − 1 bases into the run, and the count of molecules affected is stamped with the corpus. At the end of this procedure, the reads of each UMI are resolved into contigs, and the contigs of each UMI represent its true, assembled, deduplicated mRNA sequence that might be fragmented and incomplete. These contigs, grouped by the UMI and cell barcode they came from, are the direct input for model training. Supplementary Material Supp. H gives the read pipeline in full, and Supplementary Material Supp. I the invariants we test rather than assume.

### 2.2 Building the training corpus

The framework builds text over seven essential tokens that serve every organism. Four are the nucleic acid bases `A`, `C`, `G` and `T`, and one is `N` at a base the reads left ambiguous. The other two are separators, a tab (`\t`) between the contigs of one molecule and a newline (`\n`) between molecules. Models can add tokens, such as `MASK` for their own use. Under this context, the corpus holds one line per molecule carrying its contig sequences separated by tab, and a cell is a contiguous range of lines.

We provide a dedicated toolkit written in Rust, coupled with a compact binary file format for storing the corpus and loading it into training. A corpus is a set of shards, each a columnar file that holds a contiguous range of cells at one byte per token. It is up to 4.3*×* smaller than the gzipped FASTQ it was built from, depending on how duplicated the reads are, and it supports advanced features like random cell access. While loading for model training, a shard is mapped read-only into the process’s address space, so the arrays a model reads are views of mapped pages rather than copies, and only the pages a batch reads become resident in RAM. The corpus is staged to node-local storage before a run, so a page fault costs a local read rather than a round trip to a network filesystem. The data loader is also written in Rust, so it releases the global interpreter lock while its own threads work, and it puts each batch into page-locked host memory so the transfer to the device needs no staging copy. The time the training loop spends blocked on a batch therefore stays under 1% of a step. Supplementary Material Supp. D gives every corpus, and Supplementary Material Supp. E names every dataset and where it came from.

### 2.3 Architecture

In this section, we introduce the single-cell read foundation model scRM. Its trunk borrows the idea of hierarchical modeling from the byte latent transformer (Pagnoni et al., 2025), adopted for dedicating the information propagation from nucleic acid sequences to cells into five levels, namely bytes, contigs, molecules, windows and cells, where bytes, molecules and cells are the real biological entities and the contigs and windows are intermediate layers introduced in data processing. We train transformers at the byte and the molecule levels, and propagate information between levels with attention pools. The whole model we presented in this manuscript has about 1.2 million parameters, 1,182,720 specifically.

The model reads one window at a time. A window is 16,384 tokens, filled with UMIs from the same cell in random order until the next one would not fit, so it carries about 125 molecules and at most 268. Two arrays of segment identifiers travel with the tokens, one naming the contig each position belongs to and one naming its molecule, so no level boundary is written into the model. Every token is embedded into a learnable vector of width 96, and one step reads 128 cells at two windows each.

We put one transformer at each level whose elements need to be related to one another. The byte encoder is four layers of width 96 and reads the embedded tokens. Its attention is confined to one contig, so the model cannot relate bases the sequencing never observed together. By contrast, the molecule encoder, six layers of the same width, reads one vector per molecule after the hierarchical pools from bytes to contigs and then from contigs to molecules, relating every molecule in a window to every other. We apply rotary positions only for the byte encoder and make them restart at each contig, because contigs are measured sequences while a cell is an unordered set of molecules.

We apply four attention pools to propagate information through the hierarchy, once at each level boundary. Each scores the elements of a group against one learned query, turns the scores into weights, and returns the weighted sum (Ilse et al., 2018). The contig pool collapses a contig’s bytes, the molecule pool collapses a molecule’s contigs, the window pool collapses a window’s molecules and the view pool collapses a cell’s windows. All four keep the width at 96, so every pool is dimension-preserving. The four share the operation but not the parameters. The vector returned by the last of the four, the view pool over a cell’s windows, is the cell embedding, and it is what the cell-level term of the loss is computed on. Supplementary Material Supp. B gives every hyperparameter.

### 2.4 Training objective

Our objective combines three ideas from the literature. We recover masked input as in the masked autoencoder (He et al., 2022b), and we mask blocks of positions rather than single ones as in JEPA-DNA (Larey et al., 2026). Every step runs two copies of the model, a student the optimizer updates and a teacher it does not, and we hold the teacher’s weights as an exponential moving average of the student’s as in DINO (Caron et al., 2021). The student predicts the teacher’s representation of the content the student cannot see, as in I-JEPA (Assran et al., 2023), and we add a term against collapse as in LeJEPA (Balestriero and LeCun, 2025).

At the byte level *ℒ*_mlm_ recovers the masked bases. The student reads a window in which some blocks of bases have been replaced by the `MASK` token, and the teacher reads the same window unmasked. Blocks are chosen inside one contig and never over a separator, and we apply a dynamic masking ratio ranging from [0.15, 0.25] to [0.20, 0.30]. Write *x* for the window’s tokens, *h* for the student’s byte states, *K* for the masked positions and *E*_0:4_ for the embedding rows of the four bases. Cross-entropy is taken at masked positions only and over the four bases, and its head is *E*_0:4_ transposed, so it adds no parameters. Separators and ambiguous bases are never a target. *ℒ*_mlm_ is what holds the objective to the sequence, because a model that only predicted its own representations could satisfy every other term without keeping the bases at all.

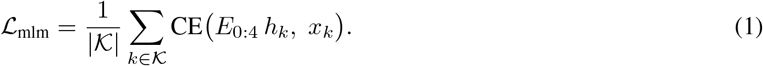

At the molecule and the cell levels we predict the teacher from the student. Write *u* for the student’s molecule vectors and *c* for its cell embedding, with 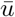 and 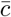 the teacher’s on the unmasked window, for the real molecule positions, *d* for the width and sg for the stop-gradient that keeps the teacher out of the backward pass. _mol_ compares *u* with 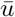 over *ℛ*, and *ℒ*_cell_ compares *c* with 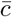. Predicting a representation rather than a reconstruction is what makes these terms JEPA, and *ℒ*_cell_ is the only one that reaches the view pool. The teacher’s momentum rises linearly from 0.996 to 1, so the target moves slowly enough for the student to predict and stops as training ends.

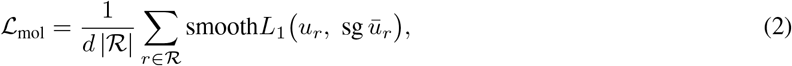

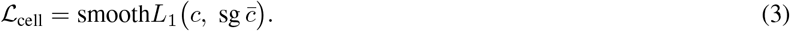

At the window level we add *ℒ*_sigreg_ against collapse, because predicting the teacher has a trivial solution: a student that maps every window to one point predicts it exactly. We use SIGReg from LeJEPA (Balestriero and LeCun, 2025), in a custom implementation. *ℒ*_reg_ tests many random one-dimensional projections of the window vectors against a standard normal, which a batch mapped to one point fails. *ℒ*_inv_ pulls a cell’s two views together. The two are mixed at *λ* = 0.0196, LeJEPA’s published value.

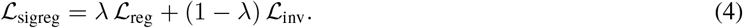

The four terms are summed with weight one each, so nothing balances them against one another.

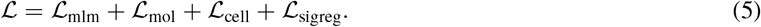

Supplementary Material Supp. C gives every term in full.

## 3 Experiments

### 3.1 Setup

#### Data

In this manuscript, we focus on 10x Genomics Chromium 3^*′*^ data, while the framework we introduced can be applied to any primer-based protocol that carries a cell barcode and a UMI, such as the 10x Chromium 5^*′*^ assay or Drop-seq. The reported model is pretrained on four mouse corpora holding 30,571 cells, 307,044,165 assembled molecules and 36.8 billion tokens, and it draws its training steps from 3,000 of those cells. Evaluation uses twelve public datasets, none of them in the corpus the reported model was trained on: five human, four mouse, one fly, one macaque and one rat sample. For all datasets, the sequencing reads and the filtered count matrix bundles are downloaded from the data provider. The cell barcode list used by scRM is obtained directly from the filtered count matrix.

#### Baselines

Four pretrained count-based single-cell foundation models are used for evaluation, all frozen and zero-shot, and each checkpointed from its authors’ own release: UCE (Rosen et al., 2026) at about 650M parameters over 36M cells in eight species, whose genes reach the model as protein embeddings from ESM2 (Lin et al., 2023) rather than through a gene vocabulary, GeneJEPA (Litman et al., 2025) at roughly 200M over 100M drug-perturbed human cells, Geneformer V2 (Theodoris et al., 2023) at 104M over 104M human cells, and scGPT (Cui et al., 2024) at 51M over 33M. We score a pretrained variational autoencoder, scVI (Lopez et al., 2018), alongside them, at 2M over 65,847 cells of a lung atlas. We fit two more count representations fresh on each evaluation dataset. The best-practice count workflow (`scanpy`) (Heumos et al., 2023; Wolf et al., 2018) is required rather than optional, because count foundation models are reported to lose to it (Kedzierska et al., 2025; Bendidi et al., 2024). A depth-only probe is the floor for every task. Its only feature is the total deduplicated UMI count, so a representation that does not beat it has learned nothing beyond how deeply that cell was sequenced.

#### Protocol

Every number we report for scRM comes from one run of the model of §2.3, trained on 3,000 mouse nervous-system cells for 8,947 steps. Nothing is fine-tuned: every representation is frozen and read zero-shot at five folds by three seeds, with the features standardized and the probe’s penalty fit inside each training fold. The estimator follows the target, not the model. Cell-line identity and cell-type annotation are read by a multinomial logistic regression whose penalty is cross-validated inside the training fold, scored in macro-F1 against a majority-class floor on the same folds; a cosine-*k*NN classifier and a small MLP read the same folds. Gene detection, pathway activity and perturbation response are read by a ridge regression whose penalty is chosen per target inside the training fold, and scored on the held-out folds as per-gene AUROC, per-module Pearson *r* and per-target AUROC. The probes do not reweight their classes; the class balance is in the metric, which averages over classes for macro-F1 and is insensitive to that balance for AUROC. AUROC is 0.5 at chance and 1 at perfect detection, and below 0.5 the ranking runs against the label; macro-F1 and accuracy run from 0 to 1. Supplementary Material Supp. F names every model scored, the checkpoint each number was read from and its size, and Supplementary Material Supp. G gives the evaluation protocols in full.

### 3.2 A scRM model of 1.2**M** parameters reaches the large count foundation models

#### A 1.2**M scRM model reaches count foundation models** 43 **to** 550 times its size

The count foundation models were trained on 33 to 104 million cells, from human or across several species. Also note that, except cell-line identity and perturbation response, the true labels of all other tasks are directly derived from count matrices. Reaching them from three thousand mouse cells shows scRM learns the species-agnostic biological signal in the reads. Table 1 puts scRM inside the range of the count foundation models on every task, and on cell-line identity it comes within 0.011 of the leading one. An exact sign test confirms those gaps are not noise: scRM is behind the best count foundation model on all 9 gene-detection datasets, all 16 pathway measurements and 23 of 26 screen targets (*p ≤* 0.0039). As a lightweight model trained on thousands of cells, we do not claim that scRM outperforms the state-of-the-art count foundation models, only that it is comparable with them.

**Table 1:**
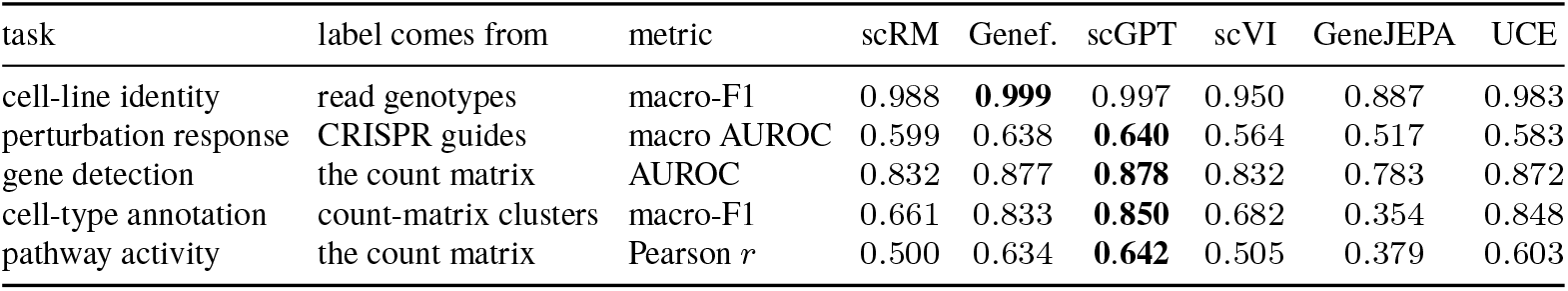
A 1.2M scRM model reaches the leading count foundation models (FMs), including scGPT, GeneJEPA, Geneformer and UCE, on all evaluation tasks. The best is Geneformer on the first row and scGPT on the rest. The metrics do not share a floor, so the rows are not comparable with each other. The scRM column is the reported model of §2.3.

#### scRM predicts biological labels well across five species

We evaluated scRM on biological label prediction. To avoid biasing our own tools, we chose the evaluation protocol published with the UCE paper rather than our own. We first tested on five human lung cancer cell lines from `sc_mixology` (Tian et al., 2019), where the ground truth labels are the cell lines themselves, assigned from the genotype mutations the sequencing observes and not from expression. The left panel of Figure 2 shows this: scRM reads 0.983 in accuracy, very close to the upper bound of the count foundation models’ own range of 0.866 to 1.000. scRM therefore assigns the correct biological label to a cell where a ground truth exists to check it against.

**Figure 2.**
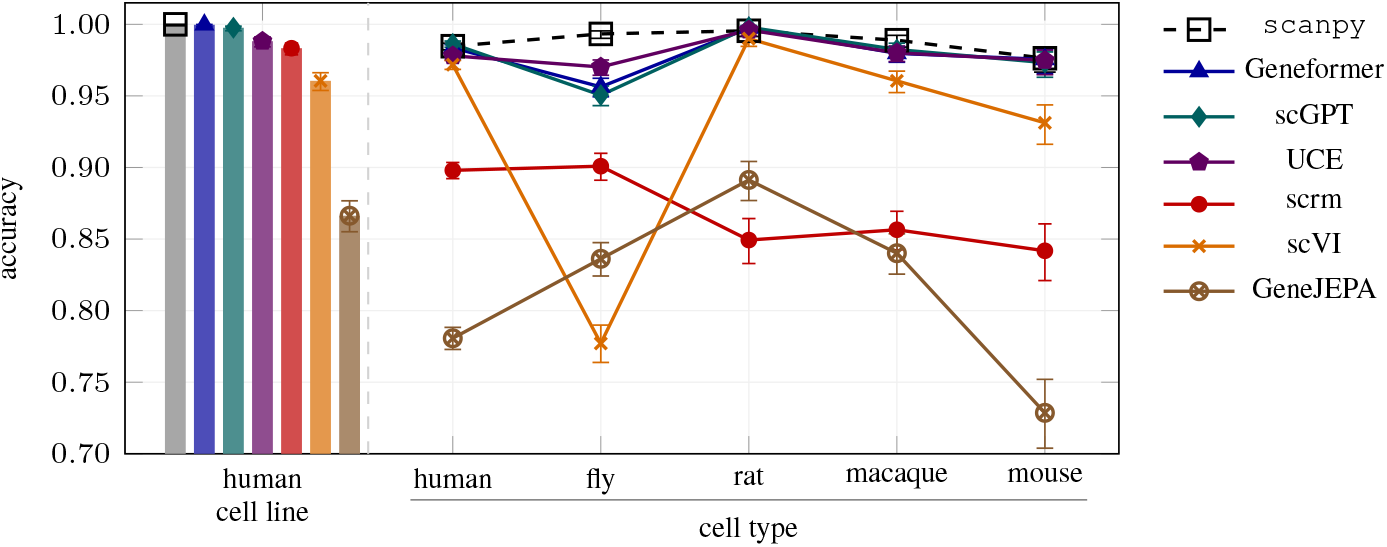
scRM trained on mouse cells predicts cell label well in all tested species, measured under UCE’s published protocol. The left panel is cell-line prediction over five human cancer cell lines labeled by demuxlet genotyping; the right panel is cell-type prediction in five species, labeled by clusters of the same count matrices. The metric is accuracy and the error bars are 95% Wilson intervals over the 1,308 to 11,026 cells each column scored. The axis starts at 0.70, so the left panel’s bars are truncated. The intervals are marginal, so overlap between two of them does not establish that the models agree; §3.2 gives the test that counts over a population.

We also applied UCE’s protocol to cell type annotation on five species: human, mouse, fly, macaque and rat. Among them the fly is the one used in the UCE paper. Note that the labels of these datasets are derived from count matrices, which disadvantages scRM because it is the only model that does not take the count matrices as the input. On the mouse panel the count foundation models reach 0.729 to 0.976 on accuracy and scRM 0.842, again inside their range. On every panel scRM reports a small gap to the best performing count-based foundation model, 0.017 on the human cell lines and 0.088 to 0.149 on the five annotation panels.

#### Models that read sequences suffer less when applied out-of-species

Every count model suffers gene name mis-matches when it meets other species, and therefore performs worse. From human to fly, scVI falls 0.195, scGPT 0.036 and Geneformer 0.027, while models that read sequences fall less or not at all, with UCE 0.008 and scRM gaining 0.003. UCE identifies a gene by its protein embedding rather than by its name, and scRM reads the bases, so neither carries a vocabulary that can fail to match. GeneJEPA also gains, 0.055 on fly, from a human baseline of 0.781 that leaves it the most room to gain. Supplementary Material Supp. K gives the ablations.

### 3.3 scRM leads where perturbation detection requires sequence-level detail

We tested perturbation detection on a perturb-seq dataset in A549 human lung carcinoma cells with 26 measurable knockdowns. The screen also declares non-targeting cells that are not perturbed, which can be used for measuring the specificity because no probe should call them active (Table 2 and Supplementary Material Supp. Table 3). We score each gene target by AUROC, where a true positive is a perturbed cell called active and a false positive is a control cell called active. Supplementary Material Supp. J gives the screen in full.

**Table 2:** AUROC on perturbation detection in a CRISPR screen, where the guide identity is from 10x Genomics feature barcoding. Every target is scored against the non-targeting control cells the screen declares. The first two rows are over all 26 targets: the macro AUROC, and the specificity in them.

| target | gene function | scRM | scanpy | Genef. | scGPT | scVI | GeneJEPA | UCE |
|---|---|---|---|---|---|---|---|---|
| overall | — | 0.599 | 0.588 | 0.638 | **0.640** | 0.564 | 0.517 | 0.583 |
| control specificity | — | 0.662 | 0.656 | **0.687** | 0.682 | 0.639 | 0.612 | 0.648 |
| *MORF4L1* | exon exclusion | **0.629** | 0.523 | 0.561 | 0.555 | 0.556 | 0.473 | 0.523 |
| *SRPK1* | exon inclusion | **0.605** | 0.439 | 0.573 | 0.564 | 0.534 | 0.530 | 0.571 |
| *ELOF1* | transcription rate | 0.920 | 0.900 | **0.945** | 0.926 | 0.746 | 0.559 | 0.804 |
| *CTNNB1* | transcription activation | 0.669 | **0.690** | 0.665 | 0.668 | 0.523 | 0.398 | 0.594 |
| *PAIP2* | translation rate | 0.555 | 0.486 | 0.571 | **0.576** | 0.507 | 0.461 | 0.566 |
| *EIF4EBP1* | translation initiation | 0.540 | 0.472 | 0.547 | 0.548 | 0.544 | 0.545 | **0.555** |

**Table 3:** Gene detection on `sc_mixology`, AUROC over the 943 Wilcoxon marker genes it can score. scRM is the reported model, trained on 3,000 mouse neuron cells, and scRM^*†*^ is the same architecture and recipe trained on `sc_-mixology`’s own cells at the same 3,000 cap. The floor column is a depth-only probe.

|  | Genef. | scGPT | scRM<sup>†</sup> | scanpy | scRM | UCE | scVI | GeneJEPA | floor |
|---|---|---|---|---|---|---|---|---|---|
| AUROC | **0.910** | 0.908 | 0.902 | 0.901 | 0.889 | 0.888 | 0.863 | 0.830 | 0.736 |

#### scRM is ahead where the perturbation changes transcript structure, indicating that it learns sequence level signals from sequencing reads

As shown in Table 2, scRM has the highest AUROC for predicting *MORF4L1* and *SRPK1*, which change transcript structure in opposite directions. H3K36me3 brings *MORF4L1* to the alternatively spliced region, and *MORF4L1* in turn draws PTB to the nascent transcript, and raising it is sufficient to force exclusion of the PTB-dependent exons (Luco et al., 2010); *SRPK1* phosphorylates the splicing factor SRSF1, and depleting it lowers that phosphorylation, holds SRSF1 out of the nucleus and so removes its ability to promote inclusion of an alternative exon (Gonçalves et al., 2014). One perturbation therefore drops an exon and the other keeps one, which is a change the overall gene count cannot express. Another piece of evidence showing that both perturbations need transcript level information is that they can be predicted by counting the hexamers in the reads. These results proved that scRM successfully learned sequence level signals and encoded them into the embedding, and that the signal is a property of the category rather than of either gene on its own.

#### Every model is ahead where the perturbation changes the gene expression profile

*ELOF1* controls transcription rate, and deleting it slows Pol II from 2.6 to 2.0 kb per minute (Geijer et al., 2021; van der Weegen et al., 2021); *CTNNB1* controls transcription activation, and TCF/LEF factors recruit it to the Wnt response elements of their target genes to turn them on (Arce et al., 2006). For this type of perturbation, scRM detects them within the range of the count-based models, which take the gene expression profile directly as their input, indicating that scRM efficiently learned the gene expression profile indirectly from the observed sequencing reads.

This study also reveals a defect of all scRNA-seq based models, because every model performs close to random guessing when the perturbation acts on translation. For example, *PAIP2* controls translation rate, displacing poly(A)-binding protein (Khaleghpour et al., 2001); *EIF4EBP1* controls translation initiation, binding eIF4E (Pause et al., 1994). Both change how much protein a transcript yields and leave the transcript in place, so no representation of RNA can carry the effect. Even UCE suffers from this defect despite taking protein embeddings as its input, because the ESM2 embeddings it takes are averaged over all the reference proteins a gene codes for, and carry no post-transcriptional or translational change from any specific cell. Single-cell ribosome profiling measures translation directly (VanInsberghe et al., 2021), so it is the modality that would carry these two perturbations. This also lays out future work on introducing multi-omics data, such as scRNA-seq alongside single-cell ATAC-seq, single-cell proteomics and spatial omics data, into single-cell raw data pretraining.

### 3.4 scRM enables dataset-specific foundation model training

Because training on the sequencing reads squeezes all juice from an scRNA-seq experiment, our framework makes it possible to train a dataset-specific single-cell foundation model. This is not open to any count foundation model, whose input is a feature count table with all low-level details discarded. We train the reported architecture on 3,000 cells of `sc_mixology` (Tian et al., 2019) — the same cell budget, step count and objective the reported model uses on its four mouse corpora, so the two models differ in which cells they read and in nothing else — and evaluate on all 3,918, showing that it achieves first-tier performance on predicting gene detection. The task is to predict whether a gene is detected in a cell, and the ground truth reads the count matrix directly, assigning every count above one to true.

The gene set to predict is `sc_mixology`’s 943 Wilcoxon marker genes differentially expressed between the Leiden clusters, obtained from its count matrix. Similar to §3.2, this disadvantages scRM because it is the only model that does not take the count matrix as the input. As shown in Table 3, the `sc_mixology`-trained scRM model reads 0.902 on AUROC, 0.013 higher than mouse cell-trained scRM at 0.889, indicating the in-domain training helps. Three seeds of this architecture, differing only in initialization, span 0.002 on this dataset and panel, so the gain is 7.8 times seed noise. By training with the specific dataset, scRM ties with the count-based models which see the answer in the first place, showing the single-cell raw data pretraining framework makes a dataset-specific model feasible.

## 4 Conclusion

In this project, we introduced the very first framework for single-cell raw data pretraining. We showed that our lightweight model trained on this framework works excellently when applied to out-of-species evaluation tasks, and reaches the performance of leading count foundation models on all five tasks, showing the advantage of training directly on single-cell reads.

These promising results also point to future directions. First, all tasks and results are still within the scope of count-matrix-based analysis, and we, the authors, did not have enough bandwidth and expertise to design tasks that can show the full potential of scRM. For example, although *MORF4L1* and *SRPK1* in the perturbation detection task shed some light on scRM’s ability to learn read-level information, that task cannot directly show that scRM learned read-level information.

Second, the scRM model we introduced is lightweight, at 1.2M parameters. It is sufficient as a showcase of our single-cell raw data pretraining framework, and one obvious future direction is to expand it to large-scale, general purpose foundation models using advanced training techniques like VISReg (Wu et al., 2026).

Third, we went to the extreme of avoiding read alignment completely and focused on scRNA-seq. We point out three directions to improve our framework. (i) The framework and its JEPA objective are not specific to scRNA-seq: they apply to other modalities such as scATAC-seq and long-read scRNA-seq, and to paired multi-omics data such as scRNA-seq with scATAC-seq or single-cell proteomics, allowing multi-modal training. (ii) Aligning reads against a pre-built transcriptome can bridge the fragmented and sparse sequence scRNA-seq captures, giving the model more transcriptional information. (iii) Even without a reference transcriptome, methods like Forseti (He et al., 2024) recover transcriptional signal from the cDNA hidden in read 1.

