## Supplementary Material for "scRM: A single-cell foundation model trained on scRNA-seq reads"

#### A Related work

**Sequence foundation models hold the other half of the problem: sequence without a cell.** This family is pretrained on reference sequence rather than on a measurement. DNA language models learn from genome sequence (Nguyen et al., 2023; Brixi et al., 2026), JEPA-DNA (Larey et al., 2026) puts a predictive objective over the same input, RNA language models move to transcript sequence (Penić et al., 2025; Chu et al., 2024; Fradkin et al., 2026). Two limitations follow from the input: a variant present in a sample and absent from the reference cannot be represented, and no individual cell appears in the input at all. Neither is a matter of capacity, so scaling these models does not reach a cell.

**Classical count methods set the bar these models must clear, and often clear it themselves.** Independent benchmarks find simple baselines matching or outperforming count foundation models on zero-shot embedding and clustering (Kedzierska et al., 2025; Bendidi et al., 2024), in-dataset annotation (Boiarsky et al., 2023), and perturbation prediction, where the training mean outperforms all seven models tested (Ahlmann-Eltze et al., 2025). A count method such as PCA or scVI (Lopez et al., 2018) is still best. So cell-level composition appears to be at its limit rather than counting being inadequate, and we therefore include count baselines throughout and do not rest our argument on composition tasks.

**Single-cell tools that do read past the count matrix each quantify one chosen phenomenon.** scQuint quantifies alternative splicing from the intron each read spans (Benegas et al., 2022), ReadZS scores how reads are distributed within genomic windows (Meyer et al., 2022), ScISOr-Seq barcodes full-length isoform reads by cell (Gupta et al., 2018), and scAPA and scDaPars locate alternative polyadenylation sites from $3'$ read coverage (Shulman and Elkon, 2019; Gao et al., 2021). Each of them aligns its reads to a reference genome first, and each returns a per-cell measurement of the one feature it was built for. scQuint and ReadZS need no isoform annotation, and they still need the genome to place a read. None of the five is pretrained, and none returns a vector that a second task can read. They are the strongest evidence that a read carries signal beyond its gene, and they keep the reference the count matrix depends on while giving up its generality.

**Two lines of work approach read-level single-cell modeling, and each stops one step short.** A cell-free RNA model pretrained on $\sim 10^{10}$ reads (Lee et al., 2025) suggests read-level pretraining scales, but it works on bulk samples with no barcode and no UMI, so the molecule and the cell do not exist in its data. SPLASH (Chaung et al., 2023) and its single-cell extension (Dehghannasiri et al., 2026) are the strongest evidence this signal is real, but they are statistical tests, with no pretraining and no transferable representation. Each ingredient exists alone, and to our knowledge scRM is the first foundation model pretrained on single-cell sequencing reads that also yields a transferable cell representation.

**Predictive objectives have reached single-cell data three times, always in count space.** This is the line of work ours belongs to. Such an objective trains a student to predict a teacher's representation of hidden content rather than to reconstruct the input, as I-JEPA (Assran et al., 2023) does for images and JEPA-DNA (Larey et al., 2026) for genomic sequence. Neither pairs that objective with recovering the input, and combining the two is ours (§2.4). GeneJEPA (Litman et al., 2025), Cell-JEPA (ElSheikh et al., 2026) and scJEPA (Zhai et al., 2026) all apply one to normalized or binned counts. GeneJEPA is therefore both our closest template and one of our baselines.

### B Hyperparameters

We read every value below from the configuration stored in the reported model's own checkpoint, so the model described here is the model we scored. Where that configuration carries a field the forward pass does not use, we name it as unused rather than report it as active.

**Vocabulary.** The vocabulary holds eight tokens and no unused ids: `A`, `C`, `G`, `T` = 0–3, `N` = 4, `TAB` = 5 between the contigs of one molecule, `NEWLINE` = 6 between molecules, and `MASK` = 7. The reconstruction head is the embedding's first four rows transposed, so it predicts over four classes and adds no parameters. We write `N` only where assembly finds two bases with equal support. We drop a read that carries an ambiguous base when the FASTQ is parsed, so `N` never enters from the input.

**Dimensions.** The byte encoder is 4 layers of width 96 with 4 heads and a SwiGLU hidden width of 256. Its rotary positions restart at every contig, with base $10^4$ and a table sized to the longest admissible molecule, 1,600 tokens, rather than to the window. The molecule encoder is 6 layers of width 96 with 4 heads and the same hidden width. Every molecule is handed position 0, which selects the identity rotation, so the molecule encoder is positionless by its input rather than by a switch. All four pools map $96 \to 96$. The configuration exposes a field that would widen the molecule pool, and we leave it unset, so the model widens nowhere. We set the normalization $\varepsilon$ to $10^{-5}$. We round the molecule slots of a batch up to a multiple of 32 so the molecule encoder's shapes stay static, and the extra slots are padding, masked everywhere and exactly zero (Supp. I). Almost all of the arithmetic one window costs is in the byte encoder, because it runs over 16,384 positions while the molecule encoder runs over the window's molecules only.

**Windows and batch.** A window is 16,384 tokens and holds whole molecules only. Molecules shorter than 60 or longer than 1,600 tokens are dropped at the corpus build, and a cell needs 500 molecules to enter a corpus. At most 268 molecules fit one window, which is the window divided by the shortest molecule and its separator. We take two views per cell and 128 cells per step, which is 256 windows per step. The reported run executed those on a single H100, which its checkpoint records as `world` = 1; the batch geometry is the same under data parallelism, and that matters because the regularizer's value depends on which rows it sees together. The packer admits three window sizes, 16,384, 32,768 and 49,152 tokens. The reported model and every arm in Supp. K use the first, and a run at another size is a separate experiment rather than a point on the same curve.

**Objective.** The objective carries four terms at weight 1 each, as §2.4 states them: reconstruction at masked positions, smooth-$L_1$ against the teacher at the molecule level and at the cell level, and the SIGReg term mixed as $\lambda \cdot \text{reg} + (1 - \lambda) \cdot \text{inv}$ with $\lambda = 0.0196$. The regularizer tests $K = 4 \times 96 = 384$ random directions, redrawn at every forward pass, in single precision. Its value depends on which window rows it sees together, so under data parallelism the rows are gathered across workers before the projector — machinery the reported single-GPU run does not exercise, and which is what makes the two placements equivalent rather than merely similar. Masking is block-structured with a block floor of 3 and at most 3 regions per contig, at a ratio drawn from $U(0.15, 0.25)$ at the start of training and from $U(0.20, 0.30)$ at its end. We build the masks in the data path. The configuration also carries weights for a count head and a molecule-contrast term, and both are zero, so neither is active.

**Projector and readout.** The projector is `Linear`(96→256) → `BatchNorm1d` → `ReLU` → `Linear` (256→256) → `BatchNorm1d` → `ReLU` → `Linear`(256→96), with no normalization on the output, and it holds 116,320 parameters in all. It exists for the regularizer and is discarded at evaluation. The readout applies the view pool to adjacent pairs of a cell's windows and takes the mean over those pairs. That mean is the only averaging between the bytes and the cell vector, and it sits outside the trained model. Every scRM number in this paper is then read from the standardized cell vectors, as Supp. G describes.

**The teacher.** The teacher is an exponential moving average of the student, with momentum scheduled linearly from 0.996 to 1.0, and it reads the window unmasked. Its pass is a forward under no gradient over the byte encoder, the molecule encoder and the pools, and only the projector has no teacher copy. Block masks and molecule positions are computed once and shared by the two passes, so the teacher adds a forward and not a second training step.

**Optimization.** We optimize with fused AdamW at $\beta = (0.9, 0.95)$ and $\varepsilon = 10^{-8}$, and we clip gradients at 1.0. The learning rate warms from $10^{-5}$ to $3 \times 10^{-4}$ over 1,000 steps and decays to $10^{-6}$. Weight decay is scheduled from 0.04 to 0.4 on the decayed group and is 0 on normalization parameters, biases and the pools' learned queries. The byte encoder runs under activation checkpointing, which is what makes the window size fit, and its attention uses a compiled block-mask builder, because the dense mask does not fit in memory at this shape.

**Size and cost.** The reported model has 1,182,720 parameters, summed from the checkpoint's state dict. We trained the reported run for 8,947 steps on 3,000 cells at seed 20260903, on a single H100 in one hour. Its log records a median of 390 ms per step, timed across the whole optimization step, which over 8,947 steps is 58 minutes. For scale, published pretraining costs in the count-model family run from four GPUs for 3 days to twenty-four for over 40 (Yuan et al., 2024; Rosen et al., 2026).

### C Objective terms in full

Let $x$ be a window's tokens and $\tilde{x}$ the masked copy the student reads. We write $h$ for the student's byte states, $u$ for its molecule states, $w$ for its window vector and $c$ for its cell vector, and barred symbols are the teacher's, computed on the *unmasked* window. $\mathcal{K}$ is the set of masked positions, $\mathcal{R}$ the real (non-padding) molecule slots, sg stop-gradient, and $E_{0:4}$ the embedding rows for A, C, G and T. All four weights are 1.0.

**Reconstruction.** The head is the embedding transposed, so it has no parameters of its own. We take the loss at masked positions only, because separators and ambiguous bases are excluded by construction:

$$\mathcal{L}_{\text{mlm}} = \frac{1}{|\mathcal{K}|} \sum_{k \in \mathcal{K}} \text{CE}\big(E_{0:4}\, h_k,\ x_k\big). \tag{1}$$

**The teacher.** The teacher is an exponential moving average of the student (Caron et al., 2021) with momentum scheduled $0.996 \to 1.0$, linearly per step as in JEPA-DNA and ending at exactly 1.0 as in I-JEPA, which freezes it as training ends. It is a copy of the student's byte encoder, molecule encoder and pools, so there is no teacher projector, and none of its parameters ever requires a gradient because the only thing that changes them is folding in the student. We use no separate predictor, and instead compare the student's own states to the teacher's at the molecule level and at the cell level, where the cell term is the only one that trains the view pool.

**The two prediction terms.**

$$\mathcal{L}_{\text{mol}} = \frac{1}{d\,|\mathcal{R}|} \sum_{r \in \mathcal{R}} \text{smooth}L_1\big(u_r,\ \text{sg}\,\bar{u}_r\big), \qquad \mathcal{L}_{\text{cell}} = \text{smooth}L_1\big(c,\ \text{sg}\,\bar{c}\big). \tag{2}$$

Both terms use the same elementwise loss, at the unit transition PyTorch takes by default:

$$\text{smooth}L_1(a,b) = \begin{cases} \frac{1}{2}(a-b)^2 & \text{if } |a-b| < 1, \\ |a-b| - \frac{1}{2} & \text{otherwise.} \end{cases} \tag{3}$$

In $\mathcal{L}_{\text{mol}}$ we sum the smooth-$L_1$ over the $d = 96$ coordinates, so the mean is over coordinates as well as slots.

**The means are taken over global counts.** We take that mean and the reconstruction mean over *global* counts rather than as a mean of per-worker means, because masked positions and real molecule slots are ragged across workers, so averaging per worker would weight workers instead of samples and would not reproduce the single-device value. $\mathcal{L}_{\text{cell}}$ needs no such correction and does not receive one, because every worker holds the same number of cells, so we take its mean over the worker's own batch.

**Anti-collapse.** The projector maps window vectors to $z \in \mathbb{R}^{N\times 96}$, and the regularizer tests $z$ against an isotropic Gaussian along $K = 4 \times 96 = 384$ random unit directions $v$, redrawn every forward pass. The reported model uses SIGReg (Balestriero and LeCun, 2025) in a custom implementation. SIGReg scores each one-dimensional slice $zv$ with the Epps–Pulley statistic, the squared distance between the slice's empirical characteristic function and the standard normal's integrated against a weight, and averages over the slices:

$$\mathcal{L}_{\text{reg}} = \frac{1}{K}\sum_{v} N \int_{0}^{t_{\max}} \left|\, \hat{\varphi}_{zv}(t) - e^{-t^2/2} \,\right|^2 w(t)\, \mathrm{d}t, \qquad t_{\max} = 3. \tag{4}$$

We evaluate the integral by the trapezoid rule at 17 points and compute $\hat{\varphi}_{zv}$ on the standardized slice.

**The mix with the invariance term.** We interpolate the regularizer with an invariance penalty $\mathcal{L}_{\text{inv}}$, the mean squared deviation of a cell's projected views from their own mean:

$$\mathcal{L}_{\text{sigreg}} = \lambda\, \mathcal{L}_{\text{reg}} + (1-\lambda)\, \mathcal{L}_{\text{inv}}, \qquad \lambda = 0.0196. \tag{5}$$

That value is LeJEPA's published $\lambda = 0.02$ written in this convex form, $0.02/1.02$.

**Precision.** The projector runs under the step's `bfloat16` autocast, and we cast its output to single precision before the regularizer, so we compute the statistic above in single precision on the projected rows.

**The gather comes before the projector.** Because the regularizer's value depends on which rows it sees together, we gather the window rows across workers *before* the projector rather than after. The projector carries batch normalization, so projecting per worker and gathering afterward is a different function from the single-device computation.

### D Corpora

**Pretraining.** We pretrain on four mouse 10x datasets, all $3'$ chemistry: `mouse_brain_5k`, `neuron_1k_v3`, `neurons_5k` and a spinal-cord dataset from an ALS model (`sod1_spinal_cord`). Their manifests record 30,571 cells, 307,044,165 molecules and 36.8 billion tokens after assembly (36,808,878,141), at 119.9 tokens per molecule.

**One token-pass.** We define one token-pass as the step count at which the tokens drawn equal that total, 8,776 steps at 256 windows of 16,384 tokens, and because every visit to a cell re-draws which of its windows are read, a pass covers most of the corpus rather than all of it. The reported model trains for 8,947 steps, so it reads 1.02 passes rather than exactly one.

**We report tokens beside the cell count.** Tokens are what the model consumes, and the cell count is not what bounds the training signal, because a cell of this corpus holds about 74 windows and a step draws two of them, so a pass over all 30,571 cells would visit each about 37 times with a different subset of its molecules each time.

**The reported model reads 3,000 of those cells.** It draws its 8,947 steps from 3,000 of them, so it visits each about 380 times.

**Availability.** All four are public 10x Genomics datasets, downloadable without registration and released under CC BY 4.0.

**The spinal-cord dataset.** It is *Mouse Spinal Cord, SOD1 Mutant vs. Wild Type Control*, run on a Chromium X, and it holds nuclei from the lumbar spinal cord of SOD1-G93A mutant mice against non-carrier littermates at 112 days, which is end-stage in that model.

**We collected no new samples.** We performed no animal procedures for any of the four. Evaluation uses previously published human data, peripheral blood from healthy donors, five immortalized cancer cell lines, a CRISPR screen in a lung carcinoma line and one targeted-genotyping dataset from a patient sample, all of it generated in prior studies under those studies' consent frameworks, with no new sample collection and no interaction with donors on our part.

**A read-level representation retains nucleotide content.** It can therefore carry germline variation that a count matrix reveals only through population-level association, so read-level corpora, and the representations trained on them, are better governed as genotype data than as expression matrices.

**The rest of the pipeline is fixed by the code and the chemistry.** The barcode whitelist, the assembly parameters and the window packer are determined by those rather than by us.

**The corpus is predominantly nuclei.** Three of the four datasets are nuclei preparations by 10x's own descriptions, and only the E18 brain-cell dataset is whole cells. The two do not present the same molecule population, since nuclear RNA carries more unspliced sequence, so what a byte model reads shifts with the preparation.

**The corpus spans more than one $3'$ kit generation.** The spinal-cord dataset uses the GEM-X Single Cell $3'$ v4 kit while the others use earlier $3'$ kits, which the pipeline handles by parsing each dataset's chemistry rather than assuming one.

**Evaluation.** Nine datasets carry the count-matrix targets, and the gene detection and pathway activity rows of Table 1 are means over them. None is in the pretraining corpus:

- **Mouse** (four, so a gap to a human-trained count model cannot be explained by species): `nuclei_2k`, `tm_10X_P7_2` (Tabula Muris, 1,645 cells, 16 expert cell-ontology classes), `heart_10k_v3`, `neuron_10k_v3`.
- **Human** (four, all cross-species for scRM and native for the count models): `sc_mixology_5cl` (3,822 labeled cells, five cancer cell lines labeled by demuxlet genotyping), `pbmc_1k_v2`, `pbmc_10k_v3`, `got_ET01` (targeted-amplicon genotypes).
- **Fly** (one, for the cross-species panel of §3): `fly_antenna`.

The perturbation screen is scored on its own task only (Table 2, Supp. J), and the macaque and rat samples appear only in the cell-label panels of Figure 2. Supp. E gives every download path.

**The two model families have mirror-image home ground.** scRM is mouse-trained, so the four human datasets are its cross-species case and it reads them with no gene mapping of any kind. The count models are human-trained and reach a mouse matrix only through a strict one-to-one ortholog bridge of 17,149 pairs, so the four mouse datasets are theirs.

**Mouse datasets are also simply easier.** On the Leiden-marker detection panel the depth-only floor, which knows nothing about species, scores 0.041 higher on them and the standard count pipeline 0.012 higher.

### E Every dataset and where it came from

Supp. Table 1 lists every dataset scRM trains on or is scored on: the twenty the reported model, the specialised model and the ablations of Supp. K require, and no others.

**How to read the last column.** Most are 10x Genomics datasets whose FASTQ download is `cf.10xgenomics.com/samples/cell-exp/RELEASE/SAMPLE/SAMPLE_fastqs.tar`, and for those the last column gives RELEASE/SAMPLE and nothing else. One row, `sod1_spinal_cord`, is a 10x dataset served from 10x's raw bucket rather than the CDN, at `s3-us-west-2.amazonaws.com/10x.files/samples/cell-exp/RELEASE/SAMPLE/`, and it is marked as such. The rest give an accession.

**Four rows are derived rather than downloaded.** The `_permuted` corpora are ablation condition C3 and have no source: each is built from the corpus above it by permuting molecule payloads across cells with every cell's molecule count, and therefore its read depth, held exactly, and no molecule left in the cell it came from. Reads are permuted, not labels, so the condition removes cell-specific sequence while preserving everything depth alone can explain.

**What *cells* counts.** It is the population the corpus was built over, after the 500-molecule floor, so it is smaller than the advertised count where a preparation is RNA-poor. A dash is a training corpus, whose population is a property of the training sampler rather than of the corpus.

Supp. Table 1: The twenty datasets this work uses. A `release/sample` entry is a 10x Genomics public dataset, downloadable at `cf.10xgenomics.com/samples/cell-exp/⟨release⟩/⟨sample⟩/`; the exception and the four derived corpora are described above. Sixteen are downloaded and four are derived. Every release was confirmed by requesting the archive itself rather than inferred from a sample name, because a sample name is not a release.

| dataset | species | cells | release/sample or accession |
|---|---|---|---|
| *Pretraining — the four the reported model draws its* 3,000 *cells from* | | | |
| mouse_brain_5k | mouse | 7,377 | 7.0.0/5k_mouse_brain_CNIK_3pv3 |
| neuron_1k_v3 | mouse | 1,301 | 3.0.0/neuron_1k_v3 |
| neurons_5k | mouse | 5,483 | 6.0.0/SC3_v3_NextGem_DI_Neurons_5K_Multiplex |
| sod1_spinal_cord | mouse | 16,410 | 10.0.0/SOD1_G93A_mouse_spinal_cord_P112_ specimen_1_Multiplex *(S3 host, not the CDN)* |
| mouse_brain_5k_permuted | mouse | 7,377 | DERIVED, not downloaded |
| neuron_1k_v3_permuted | mouse | 1,301 | DERIVED, not downloaded |
| neurons_5k_permuted | mouse | 5,483 | DERIVED, not downloaded |
| sod1_spinal_cord_permuted | mouse | 16,410 | DERIVED, not downloaded |
| *Held-out evaluation* | | | |
| pbmc_1k_v2 | human | 996 | 3.0.0/pbmc_1k_v2 |
| nuclei_2k | mouse | 2,288 | 2.1.0/nuclei_2k |
| tm_10X_P7_2 | mouse | 1,645 | s3://czb-tabula-muris-senis/10x/FASTQs/10X_P7_2 *(S3 host, not the CDN)* |
| heart_10k_v3 | mouse | 7,713 | 3.0.0/heart_10k_v3 |
| sc_mixology_5cl | human | 3,918 | SRA SRR8606521 |
| neuron_10k_v3 | mouse | 11,843 | 3.0.0/neuron_10k_v3 |
| pbmc_10k_v3 | human | 11,769 | 3.0.0/pbmc_10k_v3 |
| got_ET01 | human | 5,255 | SRA SRR7613775; genotypes GSM3309838 |
| fly_antenna | fly | 6,343 | E-MTAB-10519 (FCA43) |
| *Registered, not yet an evaluation column* | | | |
| 2500_rhesus_monkey_pbmcs_singleplex_3p_gem_x | macaque | 2,714 | 9.0.0/2500_Rhesus_Monkey_PBMCs_Singleplex_3p_ gem-x_Universal_Multiplex |
| 2500_wistar_rat_pbmcs_singleplex_3p_gem_x | rat | 2,474 | 9.0.0/2500_Wistar_Rat_PBMCs_Singleplex_3p_ gem-x_Universal_Multiplex |
| crispr_a549_5k | human | 4,867 | 6.0.0/SC3_v3_NextGem_DI_CRISPR_A549_5K_ Multiplex |

### F Every model scored, its checkpoint and its size

Every model is cited at its first mention in §3.1, so Supp. Table 2 gives the checkpoint rather than the paper, because where a release ships several sizes a comparison against a different size is a different comparison.

**The parameter counts.** The baselines' parameter counts are as published, and we sum ours from the checkpoints' own state dicts, which is why ours are exact and theirs are not.

**Our own rows are identified by recipe.** For our own rows the identifying information is the training recipe rather than a released name, so that is what the second column carries. The hyperparameters every one of them shares are in Supp. B.

**Two baselines have no checkpoint.** Two baselines are fit on each evaluation dataset rather than pretrained, so they have no checkpoint and no pretrained parameters.

**The two scRM rows.** Both scRM rows are the same architecture at the same recipe and differ only in what they read: the reported model on four mouse corpora (Supp. D), and scRM$^\dagger$ of Table 3 on `sc_mixology` alone. The second column carries the recipe they share — same parameter count, same cell budget, same step count, one corpus against

Supp. Table 2: Every model scored in this paper, the checkpoint each of its numbers was read from, its parameter count, and where that checkpoint can be downloaded.

| model | checkpoint | parameters | download |
|---|---|---|---|
| *Count baselines* | | | |
| UCE | 33-layer | ∼650M | figshare.com/articles/dataset/24320806 |
| GeneJEPA | released checkpoint | ≈170–200M | huggingface.co/elonlit/GeneJEPA |
| Geneformer | Geneformer-V2-104M | 104M | huggingface.co/ctheodoris/Geneformer |
| scGPT | whole-human | 51M | Google Drive folder 1oWh_-ZRdhtoGQ2Fw24HP41FgLoomVo-y, linked from the scGPT repository |
| scVI | Tabula Sapiens lung | 2M | huggingface.co/scvi-tools/tabula-sapiens-lung-scvi |
| scanpy | not pretrained | none | fit on each evaluation dataset |
| depth-only floor | not pretrained | 2 features | computed from the count matrix |
| *Ours* | | | |
| **scRM S-96 (reported)** | 3,000 cells, 8,947 steps | **1,182,720** | this work |
| scRM S-96 (†) | sc_mixology, 3,000 cells, 8,947 steps | 1,182,720 | this work |

four — and the cell budget is stated there because sc_mixology holds 3,918 cells and the arm was capped at the same 3,000 the reported model draws. The arms of Supp. K are described there.

**Which checkpoint each figure counts.** UCE's 650M and GeneJEPA's range are the figures those papers report, and we did not recount either from the weights we ran. UCE's release defaults to a 4-layer model and we ran the 33-layer one the paper reports, which is a manual download. Geneformer also ships a 316M model, which is its current default and which we did not score, so the 104M figure is the one we ran and it coincides with its corpus size.

### G Evaluation protocols

**Every task reads one frozen vector per cell.** Nothing is fine-tuned and no model sees a label before the probe. Every scRM artifact is embedded at or above its dataset's own window requirement, the token total of that dataset's deepest cell divided by the 16,384-token window, so reading all of a cell's reads is a property of the artifact rather than an intention. Every artifact declares a hash of its cell order that is recomputed from its own rows on read.

**The scored population is the same for every model.** It is the intersection of the cells each producer can cover, since scRM embeds a cell from its reads and covers the whole corpus while a count model can only embed a cell the matrix has a column for, in one canonical order, and we refuse an overlap below 95% outright. Each probe fits its own feature standardization and its penalty inside every training fold, at five folds by three seeds, with the folds shared across embeddings so every comparison is paired. The one quantity computed over all embedded cells rather than per fold is the scRM cell vector's own standardization, described next; it uses no labels, so it cannot leak one.

**The scRM cell vector is standardized before any probe reads it.** The artifact holds one vector per window. The cell vector is the mean over a cell's windows, and we then center and scale each of its dimensions over the cells being embedded. The raw pool output carries a shared offset so large that every pairwise cosine between cells is 1.000 to three decimals, so an angular readout on it measures nothing.

**Every baseline is read as it ships.** We do not standardize a count PCA, because its components are ordered by variance and that ordering is information the probe would lose. The baselines therefore carry the reduction `mean` and the scRM rows carry `mean_standardised`, and every report records which we applied. The ridge probes standardize their features inside each fold in any case, so the gene detection and pathway numbers do not move under this rule, while the label tasks and every angular readout do.

**Probes.** Cell-line identity and cell-type annotation are read by three probes: a multinomial logistic regression whose penalty is chosen from seven values by an inner three-fold cross-validation on the training fold, a cosine-$k$NN classifier and a small MLP trained to a fixed budget, each at five folds by three seeds, scored by macro-F1 beside a majority-class floor computed on the same folds. No probe reweights its classes and the folds are a plain random partition, so the class balance is carried by the metric rather than by the estimator: macro-F1 averages over classes and AUROC is insensitive to prevalence. The penalty is cross-validated rather than fixed because a fixed one means regularisation that drifts with the number of cells and the embedding width, which would hand the widest representation the weakest penalty and read that as representation quality. Gene detection uses ridge regression with the penalty chosen by cross-validation inside the fold, scored by out-of-fold per-gene AUROC and macro-averaged within each panel, against chance at 0.5. Pathway activity uses the same ridge and scores per-module Pearson $r$. The depth-only floor is the same probe on two features, the log of a cell's total count and the log of its genes detected. The count workflow normalizes, applies log $1p$, selects 2,000 highly variable genes, scales and reduces to 50 PCA dimensions, and the probe reads the PCA space and never the gene matrix.

**Cell-line identity.** The dataset is `sc_mixology_5cl`, the 3,822 labeled cells of a mixture of five human lung cancer cell lines (Tian et al., 2019). The label is the cell line, assigned by the dataset's authors from demuxlet genotyping, which reads variants in the reads rather than expression, so it is the one label in the suite a count matrix never observes. We score macro-F1 over five classes, and a majority-class predictor on the same folds reads 0.098.

**Perturbation response.** The task is the CRISPR screen of Supp. J, scored as AUROC per target against the screen's 388 non-targeting control cells and macro-averaged over the 26 targets with measured knockdown.

**Gene detection.** The task is whether a gene is detected in a cell, scored by AUROC over eleven panels per dataset and reported as the mean over the 99 columns the nine datasets give. We remove genes detected in fewer than 20 cells before selecting any panel.

**The eleven detection panels.** Six panels score detection as a count above zero: the top 200, 500 and 2,000 highly variable genes by `seurat_v3`, 2,000 by Pearson residuals, and two panels of per-cluster markers from a Leiden clustering of the same counts, the top 50 per cluster by Wilcoxon rank-sum under a Benjamini–Hochberg correction and the top 100 by F-statistic. Five panels score detection as a count above one, the threshold a predecessor of this evaluation used: its 500, 1,000 and 2,000 highly variable genes and its 100 F-statistic markers, and the per-cluster Wilcoxon markers at the top 100 per cluster. Both thresholds are published because the gap to the count models depends on the choice, and picking one after seeing the numbers would be choosing the metric to fit the result.

**The panel behind Table 3.** It is the per-cluster Wilcoxon panel of `sc_mixology_5cl`, scored at a count above one. It selects 976 genes, 943 of which carry a score, because the other 33 are detected in every cell of a fold or in none and leave AUROC undefined.

**Cell-type annotation.** The dataset is `tm_10X_P7_2`, 1,645 mouse cells over 16 expert cell-ontology classes from Tabula Muris. Those labels were made by clustering cells on their counts and naming each cluster from published markers, so the label depends on the count matrix at one remove. We score macro-F1, and a majority-class predictor on the same folds reads 0.023.

**Pathway activity.** The targets are fifty MSigDB hallmark module scores computed from the count matrix, in a raw and a depth-corrected variant whose nuisance is log $1p$ depth, over eight of the nine datasets. MSigDB publishes no fly collection, so we skip `fly_antenna` rather than score it through an ortholog bridge, and the row is Pearson $r$ meaned over 18 columns.

**The cross-species panels follow UCE's released protocol.** We transcribed the panels of Figure 2 and §3.2 from UCE's `fly_embedding_check` notebook: cell types with at least 100 cells, `StratifiedKFold(10)` at seed 0, `LogisticRegression(max_iter=200)`, accuracy, with their own shuffled-label column as the control. The datasets are ours, because scRM embeds reads and UCE's released files contain none. For `pbmc_10k_v3` and for the macaque and rat samples the cell types are marker-derived, a Leiden clustering of each dataset's own counts with each cluster assigned to the highest-scoring canonical immune marker set, so on those three panels the count workflow is recovering a partition its own pipeline drew and reads as an upper bound.

**The UCE numbers are our call into UCE's released code.** Before we reported any UCE number we re-embedded one of UCE's own released datasets, 35,833 mouse kidney cells, through our call into their code. The per-cell cosine to their published embedding is 0.978 on average against 0.5011 for a shuffled pairing, so a UCE number below another model's is UCE and not our invocation.

**The fly panel of Supp. Table 5 is a different task on the same dataset.** It is our three readers over the six broad classes of the 4,038 cells that carry one.

**Three of the eleven gene panels are circular by construction.** The gene sets of `markers_wilcoxon_gt1`, `markers_fstat_gt0` and `legacy_markers100` are differentially expressed between Leiden clusters computed from the same count matrix the detection targets come from, so a representation that recovers those clusters is partly predicting its own basis. The scorer records this in each panel's provenance and we keep the panels rather than dropping them, because the count models are scored on them on identical terms and because dropping a panel after seeing the numbers chooses the metric to fit the result. Table 3 is scored on the first of the three, and §3.4 says so.

**The other eight panels carry no such dependence.** They select genes by variance or by Pearson residuals, and Table 1's gene-detection row is the mean over all eleven.

**A representation whose extractor saw the evaluation cells is not a baseline.** The count workflow is fit fresh on each evaluation dataset and is the only row whose corpus is the evaluation data by construction, which every table says. scVI is the pretrained Tabula Sapiens lung checkpoint applied zero-shot and unsupervised, with no labels key and no refitting, on the same footing as the other pretrained models. The two scRM arms pretrained on `sc_mixology_-5cl`'s own reads are in-domain, so we exclude them from Table 1 and from every held-out table and report them only in Table 3.

**The same concern has a species-level form.** UCE pretrained on eight species including human, mouse and rhesus macaque (Rosen et al., 2026), so of the six panels of Figure 2 its human, mouse and macaque numbers come from species it trained on while its fly and rat numbers do not. scRM pretrained on mouse alone, so its mouse panel is the only one of the six from its own species. No evaluation dataset is in any model's training corpus, so this is about the species and not the cells.

### H The read pipeline, in full

**Being reference-free forced us to design the UMI rule rather than adopt one.** Every published rule partitions reads by gene before comparing UMIs, so inside a partition a coincidental Hamming-one neighbor is essentially impossible. We have no gene partition, so all of a cell's UMIs are compared at once and the coincidences dominate. Measured on our corpora, a UMI has 0.08 to 0.16 Hamming-one neighbors at twelve bases and 0.43 to 1.73 at ten, so collapsing connected components would chain unrelated molecules together.

**Our rule bounds group size by abundance instead.** Candidate pairs are at Hamming distance one. We direct each pair only from the rarer UMI to the strictly more abundant one, which makes the relation acyclic by construction rather than by a check. Each UMI takes its heaviest qualifying neighbor, we follow chains to a local maximum, and we emit one molecule per maximum with the reads unioned (§2.1). It differs from Smith et al. (2017) in having no equal-count tie-break and in being transitive along the count gradient.

**An over-merge costs more here than in a counting pipeline.** Because our output is sequence and not counts, an over-merge does not cost a count. It writes a chimeric contig into the corpus as though it were a real molecule.

**Assembly is per molecule, over that molecule's reads alone.** Each group is assembled with a compacted de Bruijn graph at $k = 19$ over that molecule's reads alone, built with `debruijn-rs` (Marks, 2024).

**A branch is resolved from the reads and their per-base Phred scores.** We reimplemented fgbio's `ConsensusCaller` (Fulcrum Genomics, 2026) to do it. Every observed base enters as a Phred score turned into an error probability $p$: its own base gains $\ln(1 - p)$ of log-likelihood and each of the other three gains $\ln(p/3)$, and the call is the argmax over the four with the consensus quality read off the normalized posterior. We model two error sources apart, as fgbio does, because the unit here is fgbio's unit, one UMI family. An error made before the UMI was attached is shared by every read of the molecule, so depth never clears it and a unanimous deep column caps at Q45, while an error made after is independent per read and depth does clear it.

**The first test asks whether the two candidates share one origin.** Two tests run in order. The first asks whether the candidates come from one molecule, comparing the disagreement they show against the disagreement their own Phred scores predict for a single origin, and rejecting the pair when the upper tail of a Poisson with that expectation falls below $10^{-7}$. The test is DADA2's abundance $p$-value (Callahan et al., 2016) moved from amplicon denoising to one UMI family, at a different threshold and for a stated reason: DADA2's $10^{-40}$ is calibrated for the opposite cost, where a false rejection invents a variant, so it errs toward merging, while a false merge here writes a chimera and a false split loses only the ordering between two contigs. At $10^{-40}$ two Q30 reads would need 24 disagreements in 80 columns to separate. It runs first because a consensus between two molecules is a chimera however confidently each position resolves. The predicted disagreement is the sum of the reads' own error probabilities, so the tolerance scales with their quality: over 80 columns, two Q40 reads are rejected at 4 disagreements, two Q30 reads at 6, and two Q8 reads not until 57. A flat identity cutoff cannot express that, which is why neither Velvet's 0.2 (Zerbino and Birney, 2008) nor MEGAHIT's 0.05 (Li et al., 2015) is used.

**The second test calls each disputed position.** It runs under a likelihood over the four bases from the reads covering it, writing `N` where the likelihood ties exactly. A tie is fgbio's only no-call condition, and we add a second, a called base whose consensus quality falls below Q10, because $1\times$Q31 against $1\times$Q30 is not a tie and would otherwise write a base on a posterior of 0.53. Two consecutive uncalled positions reject the pair as well, so `N` means exactly one ambiguous position and never a run. We compare positions only when the candidates line up. Equal-length candidates and non-rejoining tips do, while a rejoining insertion shifts every base after it, and there only a one-position interior may become `N`.

**A homopolymer run of at least $k$ is discarded rather than resolved.** A homopolymer run of at least $k$ collapses to a self-loop and loses its true length, so those vertices are marked used before any walk and a molecule that is nothing but the run yields no contig. The corpus stamp records the whole molecule definition as one string, `likelihood-q10-pre45-post40-omega7-walk2-consume-tip-anchored-rejoin-flank-seeded` — the consensus rule, its thresholds, and the walk that produced the contigs — and refuses to mix a corpus built under it with one built under another.

**Most molecules assemble to one contig.** Over the four pretraining corpora, 307,044,165 molecules assemble to a single contig 79.9% of the time, at a mean of 1.26 contigs per molecule and a mean contig length of 94.7 tokens, which is a read. A molecule with many contigs is not one transcript recovered deeply but several read piles that collided on one UMI. Molecules outside 60 to 1,600 tokens are dropped rather than truncated, and a cell needs 500 molecules to enter a corpus. A molecule of this corpus carries 119.9 tokens on average, so a 16,384-token window holds about 135 of them with their separators and at most 268.

**The corpus format.** A corpus is a directory of Arrow IPC shards and one manifest. A shard is a contiguous range of cells, so the ordered shard list already satisfies the row order the dataloader depends on, and every shard can be written independently. The barcode column is dictionary-encoded, because millions of rows span only thousands of cells. Arrow carries no checksum of the data itself, so we add a SHA-256 digest over each shard's tokens and one over the corpus. The corpus digest is invariant to sharding and filenames, so a rebuilt corpus can be verified against a pinned value after its predecessor is gone. Every recorded quantity is recomputed and compared when a corpus is read, so a corpus whose data no longer matches its record fails rather than trains. The manifest is written last, so a directory of shards is not yet a corpus.

**How a window is built.** A cell's molecules are cut into windows of at most 16,384 tokens, whole molecules only. No molecule can fail to fit, because a molecule is at most 1,600 tokens and the code verifies the ratio before a run starts. A view is one such window. Training draws two windows per cell and evaluation takes all of them through the same call, so the two cannot differ about what a view is. The windows partition the cell, so every molecule is in exactly one and two views never share a molecule.

**We draw views without replacement.** We rejected drawing them with replacement, because the expected overlap between two draws is the view size over the pool size, which runs from 29.8% to 0.12% across our corpora, so the relationship between a cell's two views would be a function of its library size and depth would reach the loss through the view pairing rather than through the view contents. We rejected a fixed number of molecules per view for the same kind of reason, since mean molecule length differs enough across corpora that one molecule count is 12.1k tokens on one corpus and 37.8k on another. The molecule visit order is a seeded permutation, so a cell's windows are exchangeable draws rather than its most amplified molecules first.

**How much smaller than the reads, across every corpus.** The main text quotes the best case. Measured on the sixteen corpora this work uses, each against its own gzipped FASTQ, the ratio of read size to corpus size runs from 0.96 (`heart_10k_v3`) to 4.30 (`fly_antenna`), with a median of 1.27 and a mean of 1.70. Pooled, 287 GB of reads becomes 196 GB of corpus, a ratio of 1.47. Two corpora are below one, where the corpus is the larger of the two.

**The compression is a property of duplication, and the UMI statistics say so directly.** It tracks the singleton fraction rather than the species or the depth: the two corpora that do not compress are 96.0% and 97.0% singleton UMIs, while the four that compress best — `fly_antenna`, `pbmc_1k_v2`, `nuclei_2k` and `tm_10X_P7_2`, at 2.57 to 4.30 — run 70.4% to 87.7%, with merge rates of 0.130 to 0.337 against 0.031 and 0.043. A format that stores molecules buys little on a library that is mostly singletons, because there is almost nothing to collapse.

**Two evaluation corpora have a percolating UMI graph, and we report it rather than gate on it.** The mean Hamming-1 degree of the UMI graph runs from 0.070 to 1.730 across the sixteen, and above a degree of 1.0 the graph percolates: a giant connected component forms and a directional merge rule is resolving adjacency inside it rather than between isolated families. Two corpora are above that threshold, `sc_mixology_5cl` at 1.730 and `tm_10X_P7_2` at 1.101, and both are evaluation datasets rather than pretraining ones. The assembler warns and never drops a molecule for it, and every report carries the degree and the percolation verdict per dataset. The longest merge chain anywhere is 8.

### I Invariants that are tested rather than assumed

An architecture of this shape can be wrong and still train: the loss falls, the metrics look ordinary, and the defect is in a place no scalar reports. The checks below are therefore properties of the implementation that a test asserts on every run, not observations we made once. We list them because each one caught something.

**The data-parallel step computes the same function as one device.** We assert it term by term against a single-process reference. This is not a formality. The first version of the anti-collapse term projected each worker's window rows and gathered afterward, which makes the projector's batch normalization normalize over $1/W$ of the batch, a different function, with no error and no visible symptom. The test caught it, and the fix is the gather ordering described in Appendix C.

**Every parameter receives gradient.** Two instruments run in the loop. The first reports the gradient norm of every named parameter and names any that is exactly absent or zero, which is how we find a module that is constructed but never called.

**Each loss term reaches only the parameters it should.** The second reports, for each of the four terms, its gradient norm into each parameter group. We fix the intended partition in advance, so a term whose gradient reaches a group it should not is reported as a wiring bug rather than absorbed. The cell-level term reaching the view pool and nothing else is the case this pins down.

**Padding molecule slots are exactly zero.** A window holds a variable number of molecules and the count is rounded up so the molecule encoder's shapes stay static. Those extra slots are masked at every consumer, and the invariant we assert is the stronger one, that their states are exactly zero and not merely ignored, because a masked-but-nonzero slot is the kind of defect that changes a pooled result without changing any loss.

**The compiled attention path matches a dense reference.** The training path builds its attention masks blockwise, which is required at this window size: constructing the equivalent dense mask allocates far more memory than a GPU has. The dense construction is kept as the reference implementation and the compiled path is asserted against it at small shapes on CPU, so the mask the model trains under is checked against one that can be read directly.

**Byte states at invalid positions are zeroed after the transformer, not before.** A fully masked attention row is a softmax over an empty set, whose value is undefined and differs between kernels. The implementation opens the diagonal so the kernel returns something finite and then zeroes those rows, so no pool can ever read a value that depended on which kernel ran.

### J The perturbation screen, in full

**The screen.** The screen is 10x's `SC3_v3_NextGem_DI_CRISPR_A549_5K`, A549 lung carcinoma transduced with a 93-guide pool over 46 targets plus non-targeting controls, $3'$ v3.1 chemistry. Guide calls come from the vendor's `protospacer_calls_per_cell.csv`: 4,867 cells carry a call, 3,442 exactly one, and 388 are non-targeting. Of 67 targets, 26 show real knockdown by the shipped efficiencies ($\log_2$ fold change $\leq -1$, $p < 0.05$), and only those are scored. Median 34 cells per guide, so a 46-way target call has little power and the readouts with power are per target against the 388 non-targeting cells. It is never a training corpus.

**The label cannot reach the model.** The shipped configuration declares two libraries, gene expression and Crispr Guide Capture, so guide identity sits in its own FASTQ files that the corpus build never opens.

**Searching a corpus for guide sequences does not test that.** It will mislead, because a protospacer is a 20-mer lifted out of the gene it targets, so it occurs in the transcriptome of any cell expressing that gene. The same search over a human corpus with no feature-barcode library at all returns tens of hits for exactly the pool's most expressed targets. Only the constant scaffold discriminates, and its null is a no-capture corpus rather than a uniform $4^{-20}$ expectation, which genomic sequence violates by roughly three orders of magnitude.

**Every model at every target.** Supp. Table 3 gives every representation at every one of the 26 targets, with no maximum taken anywhere. Most targets are undetectable by every representation here, including the count matrix that contains the knocked-down gene. Detectability is not predicted by knockdown strength ($r = -0.12$), cell count ($r = -0.001$), the number of differentially expressed genes ($r = +0.31$), or a shift in library size. Read on the reported checkpoint rather than the best of its seeds, scRM is above every count representation on 2 of the 26 targets, *MORF4L1* by 0.068 and *SRPK1* by 0.031. 3 seeds of the reported model spread the macro by 0.004, *MORF4L1* by 0.021 and *SRPK1* by 0.011, so only the first of those margins survives every seed.

Supp. Table 3: Every model at every target on the CRISPR screen, AUROC against the 388 non-targeting control cells, no maximum taken anywhere. This is the grid the six rows of Table 2 are drawn from, and those six targets are in bold here. $n_+$ is the number of cells carrying the target's guide. The leading score in each row is bold.

| target | $n_+$ | scRM | scanpy | Genef. | scGPT | scVI | GeneJEPA | UCE |
|---|---|---|---|---|---|---|---|---|
| *APH1A* | 204 | 0.562 | 0.645 | 0.636 | **0.677** | 0.629 | 0.602 | 0.625 |
| *BAZ1B* | 141 | 0.503 | 0.482 | 0.539 | **0.580** | 0.471 | 0.461 | 0.514 |
| *CAMLG* | 144 | 0.576 | 0.499 | 0.559 | **0.596** | 0.495 | 0.536 | 0.539 |
| *CDKN3* | 176 | 0.556 | 0.545 | **0.568** | 0.551 | 0.539 | 0.547 | 0.561 |
| ***CTNNB1*** | 63 | 0.669 | **0.690** | 0.665 | 0.668 | 0.523 | 0.398 | 0.594 |
| ***EIF4EBP1*** | 140 | 0.540 | 0.472 | 0.547 | 0.548 | 0.544 | 0.545 | **0.555** |
| ***ELOF1*** | 231 | 0.920 | 0.900 | **0.945** | 0.926 | 0.746 | 0.559 | 0.804 |
| *EZH2* | 183 | 0.599 | **0.865** | 0.802 | 0.816 | 0.669 | 0.547 | 0.676 |
| *GSK3A* | 255 | 0.594 | 0.598 | 0.661 | **0.678** | 0.605 | 0.528 | 0.595 |
| *H2AFY* | 142 | 0.555 | 0.483 | **0.588** | 0.561 | 0.499 | 0.491 | 0.498 |
| *HDAC1* | 122 | 0.564 | 0.474 | 0.606 | **0.653** | 0.509 | 0.523 | 0.624 |
| *HRAS* | 97 | 0.571 | 0.530 | 0.580 | **0.614** | 0.556 | 0.503 | 0.577 |
| *JUN* | 227 | 0.630 | 0.591 | **0.716** | 0.625 | 0.610 | 0.537 | 0.598 |
| *KDM1A* | 67 | 0.659 | 0.772 | **0.805** | 0.769 | 0.697 | 0.548 | 0.718 |
| *LMNA* | 69 | 0.578 | 0.448 | **0.651** | 0.634 | 0.502 | 0.573 | 0.576 |
| ***MORF4L1*** | 113 | **0.629** | 0.523 | 0.561 | 0.555 | 0.556 | 0.473 | 0.523 |
| ***PAIP2*** | 178 | 0.555 | 0.486 | 0.571 | **0.576** | 0.507 | 0.461 | 0.566 |
| *PPIB* | 210 | 0.624 | 0.568 | **0.788** | 0.637 | 0.591 | 0.551 | 0.577 |
| *PPP2CA* | 58 | 0.601 | **0.702** | 0.667 | 0.656 | 0.658 | 0.466 | 0.568 |
| *RAB1A* | 86 | 0.515 | 0.501 | 0.470 | **0.532** | 0.404 | 0.454 | 0.525 |
| *SET* | 102 | 0.626 | 0.596 | 0.563 | **0.637** | 0.603 | 0.550 | 0.573 |
| *SMARCD1* | 160 | 0.535 | 0.699 | 0.690 | **0.702** | 0.501 | 0.514 | 0.539 |
| ***SRPK1*** | 143 | **0.605** | 0.439 | 0.573 | 0.564 | 0.534 | 0.530 | 0.571 |
| *SUZ12* | 136 | 0.592 | 0.614 | 0.604 | **0.617** | 0.588 | 0.520 | 0.547 |
| *UBE2N* | 95 | 0.576 | 0.566 | 0.598 | **0.620** | 0.533 | 0.506 | 0.573 |
| *UBXN1* | 77 | 0.651 | 0.606 | 0.640 | **0.654** | 0.596 | 0.507 | 0.550 |
| macro | — | 0.599 | 0.588 | 0.638 | **0.640** | 0.564 | 0.517 | 0.583 |

**The explicit sequence-feature probes.** Each is an explicit per-cell matrix over the same cells, scored through the same protocol as the embeddings. Junction usage, standing for splicing, reaches 0.5994 macro and passes 0.60 on twelve of the twenty-six targets. A polyadenylation index reaches 0.4859 from annotated $3'$ ends and 0.5066 built de novo, each passing 0.60 on one target. All three are below a plain coverage matrix over the same genes, which reaches 0.6756 and passes 0.60 on nineteen, and a permuted null reads 0.4661.

**The non-targeting controls.** The screen declares one control category, 388 cells carrying guides with no genomic target site, and its feature reference names them outright. It flags no positive control, so the controls give a floor and no ceiling.

**First they define a clean negative class.** Table 2 scores every target against these controls, which asks whether a cell is perturbed at all. Scoring a target against the remaining guide-bearing cells instead asks a different question – whether a perturbation is separable from a mixture of the other twenty-five, every one of them perturbed – and that mixture is not a negative class the screen declares.

**Second they measure what nothing scores.** Splitting the controls in half at random and asking the identical probe to separate the halves must return 0.5, because the split is invented and every cell is unperturbed. It returns 0.46 to 0.49 instead, and the value differs by model, so it is not a constant offset. The ridge fits the invented labels on the training folds and meets unseen cells on the test fold, which at this sample size reads as slight anti-prediction. Assuming 0.50 was therefore understating every model. A second floor agrees with the first to within $\pm 0.003$ for every model, obtained by permuting the labels of the real mixture with its composition preserved, which rules out a depth or batch artefact separating perturbed from unperturbed cells in general.

### K Ablations

The conditions of this study test the explanations for Table 1 other than pretraining on reads. Each differs from the reported model in one respect: three random initializations of the architecture with no training (C2), an arm trained on the same corpora with every molecule moved to a random cell (C3), a two-number depth readout (C4), a table of hexamer frequencies with nothing trained (C5), and the same recipe with the attention pools replaced by means (C7).

**The instrument.** We score every condition on one instrument, gene detection AUROC over the `markers_wilcoxon_gt1` panel, meaned over the 9 evaluation datasets, because every condition moves one factor and the panel is not one of them. The panel is the one Table 1 reports gene detection on, so a condition's level and that table's detection row are the same measurement read at different precision. As shown in Supp. Table 4, the reported model reads $0.8319$ on AUROC on this instrument.

**When a difference counts.** We establish a difference when it clears the seed floor and has the same sign on all 9 datasets, which a two-sided sign test puts at $p = 1/256$, and a difference inside the floor is null whatever its sign pattern. We take the floor, $0.0024$ on AUROC, from 3 seeds of the reported model's recipe, which differ in nothing but the initialization and are scored on all 9 datasets, because what a delta has to exceed is the reported model's own reseed spread rather than a threshold we choose. Their mean range is $0.00234$ on AUROC, which the floor rounds up, and their widest single dataset ranges $0.0060$ on AUROC, wider than the floor because a floor on the mean is not a bound on one dataset.

**One condition cannot be parameter-matched, and it is not the objective.** Every condition but C7 is built on the reported model's own architecture, so the untrained initializations of C2 each hold $1{,}182{,}720$ parameters and the same architecture hash as the reported model, and their $\Delta$ carries no width difference at all. C7 cannot be matched that way, because a mean pool has no parameters, so replacing the window and view pools and replacing all four remove $18{,}624$ ($1.6\%$) and $37{,}248$ ($3.1\%$) of them. There is therefore no parameter-matched form of C7's question, and its $\Delta$ is read with that in mind rather than against a matched control that does not exist. We print $\Delta$ beside every verdict.

Supp. Table 4: Every condition of the study on gene detection AUROC over the `markers_wilcoxon_gt1` panel, meaned over the 9 evaluation datasets. $\Delta$ is against the reported model and $n/9$ is the number of datasets on which the condition is ahead of it. A condition is established when $|\Delta|$ clears the $0.0024$ floor and $n/9$ is 0 or 9. The reported model's cell vectors have an effective rank of 11.5 of 96 over the same datasets.

| | condition | what it removes | AUROC | $\Delta$ | $n/9$ | verdict |
|---|---|---|---|---|---|---|
| C1 | reported model | | 0.8319 | — | — | reference |
| C2 | untrained, init 1 | the architecture and the pooling do it, not the pretraining | 0.8220 | −0.0099 | 0/9 | established |
| | untrained, init 2 | | 0.8241 | −0.0078 | 0/9 | established |
| | untrained, init 3 | | 0.8214 | −0.0105 | 1/9 | clears floor, mixed |
| C3 | molecules permuted across cells, depth held | it is not cell-specific sequence at all | 0.8291 | −0.0027 | 4/9 | clears floor, mixed |
| C4 | depth only | it is library size | 0.7455 | −0.0864 | 0/9 | established |
| C5 | hexamer frequencies, nothing trained | no learning is needed | 0.8477 | +0.0158 | 7/9 | clears floor, mixed |
| C7 | window and view pools as means | the learned pooling is not load-bearing | 0.8312 | −0.0007 | 1/9 | null |
| | all four pools as means | | 0.8295 | −0.0024 | 3/9 | clears floor, mixed |

**C2, pretraining against an untrained copy of the architecture.** We take the reported model's architecture at three random initializations, read it through the same pools, and never train it, so that the condition holds the architecture and the readout fixed and removes the pretraining alone. As shown in Supp. Table 4, the untrained initializations read $0.8220$, $0.8241$ and $0.8214$ on AUROC, which are $-0.0099$, $-0.0078$ and $-0.0105$ against the reported model on the same instrument. They are ahead of it on 0, 0 and 1 of the 9 datasets, so the verdicts are established, established and clears floor, mixed. They spread $0.0027$ on AUROC across each other, which is outside the $0.0024$ floor.

**C3, an arm trained on permuted molecules.** We train C3 on permuted twins of the four pretraining corpora, in which every molecule's sequence moves to a random cell and every cell keeps its molecule count, so depth is held exactly and no molecule stays in its own cell. As shown in Supp. Table 4, it reads $0.8291$ on AUROC, $-0.0027$ against the reported model, and it is ahead of the reported model on $4$ of the $9$ datasets, so under the floor rule above the verdict is clears floor, mixed.

**C4, the depth readout.** The depth readout gives two numbers per cell, the log of its total count and the log of its genes detected, and it enters the study because library size is the explanation a detection metric is most exposed to. As shown in Supp. Table 4, it reads $0.7455$ on AUROC, $-0.0864$ against the reported model, and it is ahead of the reported model on $0$ of the $9$ datasets, so the verdict is established. The dataset spread of a member is the difference between its ninth and first decile across the $9$ datasets, and for the reported model it is $0.1537$ on AUROC, larger than the widest $|\Delta|$ of Supp. Table 4 at $0.0864$, so the sign count stands beside every mean.

**Usable dimensions.** We count *usable dimensions* as the exponential of the entropy of the cell vectors' normalized singular values, which is the number of directions the vectors vary along. On the full embeddings of the $9$ evaluation datasets the reported model's cell vectors use $11.5$ of their $96$ dimensions, the count pipeline's PCA $44.0$ of $50$, the hexamer table $26.8$ of $50$ and Geneformer $413.0$ of $768$.

**C7, the attention pools replaced by means.** We replace the attention pools with plain means, in one arm the window and view pools that build the cell vector and in the other all four, so that the condition moves the pooling alone. As shown in Supp. Table 4, the window and view means read $0.8312$ on AUROC and all four means $0.8295$, which are $-0.0007$ and $-0.0024$ against the reported model. The arms are ahead of it on $1$ and $3$ of the $9$ datasets, so the verdict is null for the first arm and clears floor, mixed for the second. The two arms differ from each other by $0.0017$ on AUROC.

**C5, a table of hexamer frequencies with nothing trained.** C5 counts, for every cell, the frequency of each of the $4^6 = 4{,}096$ hexamers over the same corpus tokens the model reads, never crossing an `N` or a separator, takes the log of one plus the scaled frequency, and reduces the table to its first $50$ principal components. Nothing is trained, and the result has $50$ dimensions against the model's $96$. As shown in Supp. Table 4, on the ablation instrument it reads $0.8477$ on AUROC, $+0.0158$ against the reported model, and it is ahead of it on $7$ of the $9$ datasets, so the verdict is clears floor, mixed. The reported model is ahead on the remaining $2$, by $0.0349$ and $0.0010$ on AUROC. On the two targets of §3.3 it reads $0.708$ on *MORF4L1* against $0.629$ and $0.626$ on *SRPK1* against $0.605$, reading the corpus and nothing else.

Supp. Table 5: The reported model beside every count foundation model on every task the main text scores. Cell-line identity, cell-type annotation and the fly panel are macro-F1 under the linear reader of Supp. G; gene detection is the AUROC of Table 1 over the `markers_wilcoxon_gt1` panel; pathway activity is Pearson $r$; perturbation response is the macro AUROC of Table 2. Every count foundation model of Table 1 is a column, rather than the best of them.

| task, dataset | reader or metric | scRM | Genef. | scGPT | scVI | GeneJEPA | UCE |
|---|---|---|---|---|---|---|---|
| cell-line identity | linear | 0.9881 | 0.9993 | 0.9969 | 0.9499 | 0.8869 | 0.9833 |
| cell-type annotation, mouse | linear | 0.6608 | 0.8329 | 0.8499 | 0.6823 | 0.3544 | 0.8481 |
| cell-type annotation, fly | linear | 0.4961 | 0.5639 | 0.5742 | 0.2934 | 0.3908 | 0.6756 |
| gene detection, 10 datasets | AUROC, 10 columns | 0.8283 | 0.8747 | 0.8757 | 0.8300 | 0.7830 | 0.8672 |
| pathway activity, 9 datasets | Pearson $r$, 18 columns | 0.4940 | 0.6305 | 0.6390 | 0.5024 | 0.3724 | 0.5957 |
| perturbation response, the CRISPR screen | macro AUROC | 0.5994 | 0.6382 | 0.6402 | 0.5640 | 0.5165 | 0.5834 |

**What the hexamer table is.** The hexamer table reads the same bytes the model reads and uses no reference, no annotation and no gene vocabulary, so it is a read-level representation and not a count baseline. It is the only entry in this paper that differs from the reported model in exactly one respect, that nothing was trained. Every label in Supp. Table 5 except cell-line identity is derived from a count matrix. Table 1 carries no hexamer column, and this appendix is where we report it.

Mahmoud Assran, Quentin Duval, Ishan Misra, Piotr Bojanowski, Pascal Vincent, Michael Rabbat, Yann LeCun, and Nicolas Ballas. Self-supervised learning from images with a joint-embedding predictive architecture. In *IEEE/CVF Conference on Computer Vision and Pattern Recognition (CVPR)*, 2023.

Randall Balestriero and Yann LeCun. LeJEPA: Provable and scalable self-supervised learning without the heuristics. *arXiv preprint arXiv:2511.08544*, 2025.

Ihab Bendidi, Shawn Whitfield, Kian Kenyon-Dean, Bram Schuurmans, Emmanuelle Denton, et al. Benchmarking transcriptomics foundation models for perturbation analysis: one PCA still rules them all. *arXiv preprint arXiv:2410.13956*, 2024.

Gonzalo Benegas, Jonathan Fischer, and Yun S. Song. Robust and annotation-free analysis of alternative splicing across diverse cell types in mice. *eLife*, 11:e73520, 2022. doi: 10.7554/eLife.73520.

Rebecca Boiarsky, Nalini M. Singh, Alejandro Buendia, Gad Getz, and David Sontag. A deep dive into single-cell RNA sequencing foundation models. *bioRxiv*, 2023. doi: 10.1101/2023.10.19.563100.

Garyk Brixi, Matthew G. Durrant, Jason Ku, Michael Poli, et al. Genome modelling and design across all domains of life with Evo 2. *Nature*, 2026. doi: 10.1038/s41586-026-10176-5.

Benjamin J. Callahan, Paul J. McMurdie, Michael J. Rosen, Andrew W. Han, Amy Jo A. Johnson, and Susan P. Holmes. DADA2: high-resolution sample inference from Illumina amplicon data. *Nature Methods*, 13(7):581–583, 2016. doi: 10.1038/nmeth.3869.

Mathilde Caron, Hugo Touvron, Ishan Misra, Hervé Jégou, Julien Mairal, Piotr Bojanowski, and Armand Joulin. Emerging properties in self-supervised vision transformers. In *IEEE/CVF International Conference on Computer Vision (ICCV)*, 2021.

Kaitlin Chaung, Tavor Z. Baharav, Geoffrey Henderson, Ivan N. Zheludev, Peter L. Wang, and Julia Salzman. SPLASH: A statistical, reference-free genomic algorithm unifies biological discovery. *Cell*, 186(25):5440–5456, 2023. doi: 10.1016/j.cell.2023.10.028.

Yanyi Chu, Dan Yu, Yupeng Li, et al. A 5' UTR language model for decoding untranslated regions of mRNA and function predictions. *Nature Machine Intelligence*, 6:449–460, 2024. doi: 10.1038/s42256-024-00823-9.

Roozbeh Dehghannasiri, Marek Kokot, Alexander L. Starr, Jamie Maziarz, Tal Gordon, Serena Y. Tan, Peter L. Wang, Ayelet Voskoboynik, Jacob M. Musser, Sebastian Deorowicz, and Julia Salzman. Reference-free discovery with barcoded single-cell sequencing. *Nature Biotechnology*, 2026. doi: 10.1038/s41587-026-03084-6.

Ali ElSheikh, Rui-Xi Wang, Weimin Wu, Yibo Wen, Payam Dibaeinia, Jennifer Yuntong Zhang, Jerry Yao-Chieh Hu, Mei Knudson, Sudarshan Babu, Shao-Hua Sun, Aly A. Khan, and Han Liu. Cell-JEPA: Latent representation learning for single-cell transcriptomics. *arXiv preprint arXiv:2602.02093*, 2026. count-space JEPA on an scGPT backbone; no public code as of this writing.

Philip Fradkin et al. Orthrus: toward evolutionary and functional RNA foundation models. *Nature Methods*, 23: 935–945, 2026. doi: 10.1038/s41592-026-03064-3.

Fulcrum Genomics. fgbio: tools for working with genomic and high throughput sequencing data, 2026. URL `https://doi.org/10.5281/zenodo.10456900`. Software, `github.com/fulcrumgenomics/fgbio`, archived on Zenodo; the per-base consensus model of `ConsensusCaller.scala`.

Yipeng Gao, Lei Li, Christopher I. Amos, and Wei Li. A single-cell RNA-seq based computational method to infer alternative polyadenylation events. *Genome Research*, 31(10):1856–1866, 2021. doi: 10.1101/gr.271346.120. scDaPars.

Ishaan Gupta et al. Single-cell isoform RNA sequencing characterizes isoforms in thousands of cerebellar cells. *Nature Biotechnology*, 36(12):1197–1202, 2018. doi: 10.1038/nbt.4259.

Kasia Z. Kedzierska, Lorin Crawford, Ava P. Amini, and Alex X. Lu. Zero-shot evaluation reveals limitations of single-cell foundation models. *Genome Biology*, 26:101, 2025. doi: 10.1186/s13059-025-03574-x.

Ariel Larey, Eyal Dahan, Avi Bleiweiss, Ron Kellerman, Gil Leib, Omri Nayshool, Dan Ofer, Tal Zinger, Dan Dominissini, Gideon Rechavi, Nicole Bussola, Seungwoo Lee, Shane O'Connell, Duy Hoang, Matthias Wirth, Alexander W. Charney, Yonatan Shavit, and Noa Daniel. JEPA-DNA: Grounding genomic foundation models through joint-embedding predictive architectures. *arXiv preprint arXiv:2602.17162*, 2026.

Sang Lee, Ryan Kim, and Beomsoo Kim. Beyond annotation: Leveraging raw RNA-seq reads via foundation models for multi-cancer early detection. *medRxiv*, 2025. doi: 10.64898/2025.12.17.25342371. Posted 2025-12-18; bulk cfRNA, not single-cell.

Dinghua Li, Chi-Man Liu, Ruibang Luo, Kunihiko Sadakane, and Tak-Wah Lam. MEGAHIT: an ultra-fast single-node solution for large and complex metagenomics assembly via succinct de Bruijn graph. *Bioinformatics*, 31(10): 1674–1676, 2015. doi: 10.1093/bioinformatics/btv033.

Elon Litman, Tyler Myers, Vikash Agarwal, Arjun Gopinath, Oliver Li, Eshan Mittal, and Timothy Kassis. GeneJEPA: A predictive world model of the transcriptome. *bioRxiv*, 2025. doi: 10.1101/2025.10.14.682378.

Romain Lopez, Jeffrey Regier, Michael B. Cole, Michael I. Jordan, and Nir Yosef. Deep generative modeling for single-cell transcriptomics. *Nature Methods*, 15:1053–1058, 2018. doi: 10.1038/s41592-018-0229-2.

Patrick Marks. debruijn: de Bruijn graph construction and compaction for DNA sequences, 2024. URL `https://github.com/10XGenomics/rust-debruijn`. Software, version 0.3.4.

Elisabeth Meyer, Kaitlin Chaung, Roozbeh Dehghannasiri, and Julia Salzman. ReadZS detects cell type-specific and developmentally regulated RNA processing programs in single-cell RNA-seq. *Genome Biology*, 23:226, 2022. doi: 10.1186/s13059-022-02795-8.

Eric Nguyen, Michael Poli, Marjan Faizi, Armin Thomas, Callum Birch-Sykes, Michael Wornow, Aman Patel, Clayton Rabideau, Stefano Massaroli, Yoshua Bengio, Stefano Ermon, Stephen A. Baccus, and Christopher Ré. HyenaDNA: Long-range genomic sequence modeling at single nucleotide resolution. In *Advances in Neural Information Processing Systems (NeurIPS)*, 2023. arXiv:2306.15794.

Rafael Josip Penić et al. RiNALMo: general-purpose RNA language models can generalize well on structure prediction tasks. *Nature Communications*, 16:5671, 2025. doi: 10.1038/s41467-025-60872-5.

Yanay Rosen, Yusuf Roohani, Ayush Agarwal, Leon Samotorcan, Stephen R. Quake, and Jure Leskovec. Universal cell embedding provides a foundation model for cell biology. *Nature*, 2026. doi: 10.1038/s41586-026-10689-z.

Eldad David Shulman and Ran Elkon. Cell-type-specific analysis of alternative polyadenylation using single-cell transcriptomics data. *Nucleic Acids Research*, 47(19):10027–10039, 2019. doi: 10.1093/nar/gkz781.

Tom Smith, Andreas Heger, and Ian Sudbery. UMI-tools: modeling sequencing errors in unique molecular identifiers to improve quantification accuracy. *Genome Research*, 27:491–499, 2017. directional adjacency.

Luyi Tian, Xueyi Dong, Saskia Freytag, Kim-Anh Lê Cao, Shian Su, Abolfazl JalalAbadi, Daniela Amann-Zalcenstein, Tom S. Weber, Azadeh Seidi, Jafar S. Jabbari, Shalin H. Naik, and Matthew E. Ritchie. Benchmarking single cell RNA-sequencing analysis pipelines using mixture control experiments. *Nature Methods*, 16:479–487, 2019. doi: 10.1038/s41592-019-0425-8. the sc_mixology benchmark.

Xinyu Yuan, Zhihao Zhan, Zuobai Zhang, Manqi Zhou, Jianan Zhao, Boyu Han, Yue Li, and Jian Tang. Cell-ontology guided transcriptome foundation model. In *Advances in Neural Information Processing Systems (NeurIPS)*, 2024. arXiv:2408.12373.

Daniel R. Zerbino and Ewan Birney. Velvet: algorithms for de novo short read assembly using de Bruijn graphs. *Genome Research*, 18(5):821–829, 2008. doi: 10.1101/gr.074492.107.

Yuyao Zhai, Weixu Wang, Till Richter, and Fabian J. Theis. scJEPA: Dual-space self-supervised learning for single-cell representations. In *LMRL Workshop at the International Conference on Learning Representations (ICLR)*, 2026. denoising in expression space + JEPA latent prediction + SIGReg; no public code as of this writing.
